# Delay of human early development via in vitro diapause

**DOI:** 10.1101/2023.05.29.541316

**Authors:** Dhanur P. Iyer, Vera A. van der Weijden, Heidar Heidari Khoei, Afshan McCarthy, Teresa Rayon, Claire S. Simon, Ilona Dunkel, Sissy E. Wamaitha, Kay Elder, Phil Snell, Leila Christie, Edda G. Schulz, Kathy K. Niakan, Nicolas Rivron, Aydan Bulut-Karslioglu

## Abstract

Many mammals can control the timing of gestation and birth by pausing embryonic development at the blastocyst stage. It is unknown whether the capacity to pause development is conserved, in general across mammals, and more specifically in humans. Activity of the growth regulating mTOR pathway governs developmental pausing in the mouse (*1*). Here we show a stage-specific capacity to delay the progression of human development via mTOR inhibition. In this context, human blastoids and pluripotent stem cells in naïve and naïve-like, but not primed, states can be induced to enter a dormant state, which is reversible at the functional and molecular level. Comparative analysis of mouse and human naïve cells’ longitudinal response to mTORi revealed distinct temporal dynamics and metabolic requirements of dormancy in each species. Mouse and human blastocysts show similar tissue-specific patterns of mTOR pathway activity, suggesting that the mTOR pathway may be a conserved regulator of blastocyst development and timing in both species. Our results raise the possibility that the developmental timing of the human embryo may be controllable, with implications for reproductive therapies.

## INTRODUCTION

Timing of early mammalian development can be controlled in vivo via natural preservation of the blastocyst stage embryo for weeks to months in a dormant state (*2, 3*). This process, termed embryonic diapause or developmental pausing, equips over 130 species of mammals with a strategy to optimize the timing of birth and minimize the impact of environmental stressors on progeny. In diapause, the embryo shows minimal anabolic activity, coupled with reduced or diminished proliferation (*3, 4*). In the mouse, naïve pluripotency characteristics are retained during diapause, indicating that this primordial state can be stabilized and maintained in the absence of proliferation (*5*).

Although not all mammals employ diapause in their reproductive cycle, interspecies uterine transfer experiments suggest that the capacity to undergo diapause may be present across mammals (*6, 7*). Yet, significant species-specific differences in regulatory networks and morphological organization exist as well (*8–12*). Detailed analysis in recent years revealed conserved as well as distinct signaling pathways utilized by humans, mice, and other species to determine cell fates in the early embryo (*13–15*). Given these differences, whether mammals harbor conserved regulatory networks to enable diapause is unclear.

The possibility of diapause in humans has been raised, but a convincing case has never been shown (*16*). The large variation in the implantation window and developmental capacity of human embryos, as well as hormonal response and uterine receptiveness of the mothers, precludes studying human diapause in vivo beyond case studies that provide circumstantial evidence (*17*). However, surplus in vitro fertilized embryos, embryonic stem cells (ESCs) derived from early embryos, cells reprogrammed to generate induced pluripotent stem cells (iPSCs), or embryo-like models generated from PSCs can be used to test diapause potential. Few attempts to prolong the culture duration of human embryos in culture failed to retain morphology or provide sufficient evidence (*18, 19*). Importantly, the reversibility of pausing needs to be documented functionally and/or molecularly in order to make a substantiated case of human diapause.

We have previously discovered that the mTOR pathway governs developmental pausing in mice (*1*). Inhibition of mTOR (mTORi), and therefore cellular growth, induces a diapause-like state in both mouse blastocysts and ESCs. Mouse blastocysts can be sustained in vitro for several weeks under mTORi-induced pausing, while ESCs can be constantly maintained under mTORi. This paused pluripotent state is reversible and pause-released embryos and ESCs can give rise to live, fertile mice and high-grade chimeras, respectively (*1*). Downstream of mTORi, paused cells display reduced global transcription and translation, and altered metabolic networks (*20*). The transcriptional profile of mTORi-paused ESCs closely resembles that of the in vivo diapaused epiblast, suggesting that mTOR may be the master regulator of developmental pausing also in vivo.

Here we show that human model embryos (blastoids) as well as pluripotent stem cells (PSCs) can be put into a diapause-like, stable, and reversible dormant state via mTORi. By comparing naïve (PXGL), naïve-like (RSeT) and primed (mTeSR) culture conditions, we show a stage-specific diapause response in human PSCs that shares common features with mouse diapause. We also find species-specific pathway usage and a distinct dormancy progression in human cells. These results raise the possibility of modulating the timing of early human development by extending the time window of developmental competence at the pre-implantation stage.

## RESULTS

### Similar pattern and sensing of mTOR pathway activity in mouse and human embryos

Since the mTOR pathway is a major controller of the dormancy vs proliferation decision in the mouse blastocyst, we first characterized mTOR activity levels and pattern in human blastocysts in comparison to the mouse (Figure 1). The prominent mTOR downstream target phospho-S6 (pS6) showed a highly similar pattern in both species, with polar trophectoderm (TE) showing higher activity than both mural TE and the inner cell mass (ICM) (Figure 1a-b). The TE showed a salt-and-pepper pS6 pattern in both species, especially at the mural side. The same pattern was also observed for other mTOR targets in mouse embryos, indicating that the pattern does not only reflect protein synthesis but mTOR activity in general (Figure S1a). Of note, dormancy progresses from the mural to the polar side in natural mouse diapause (*21*). The weaker but consistent staining of mTOR targets in the ICM suggested that the mTOR pathway is active in pluripotent cells, albeit at lower levels compared to the polar TE.

**Figure 1.**
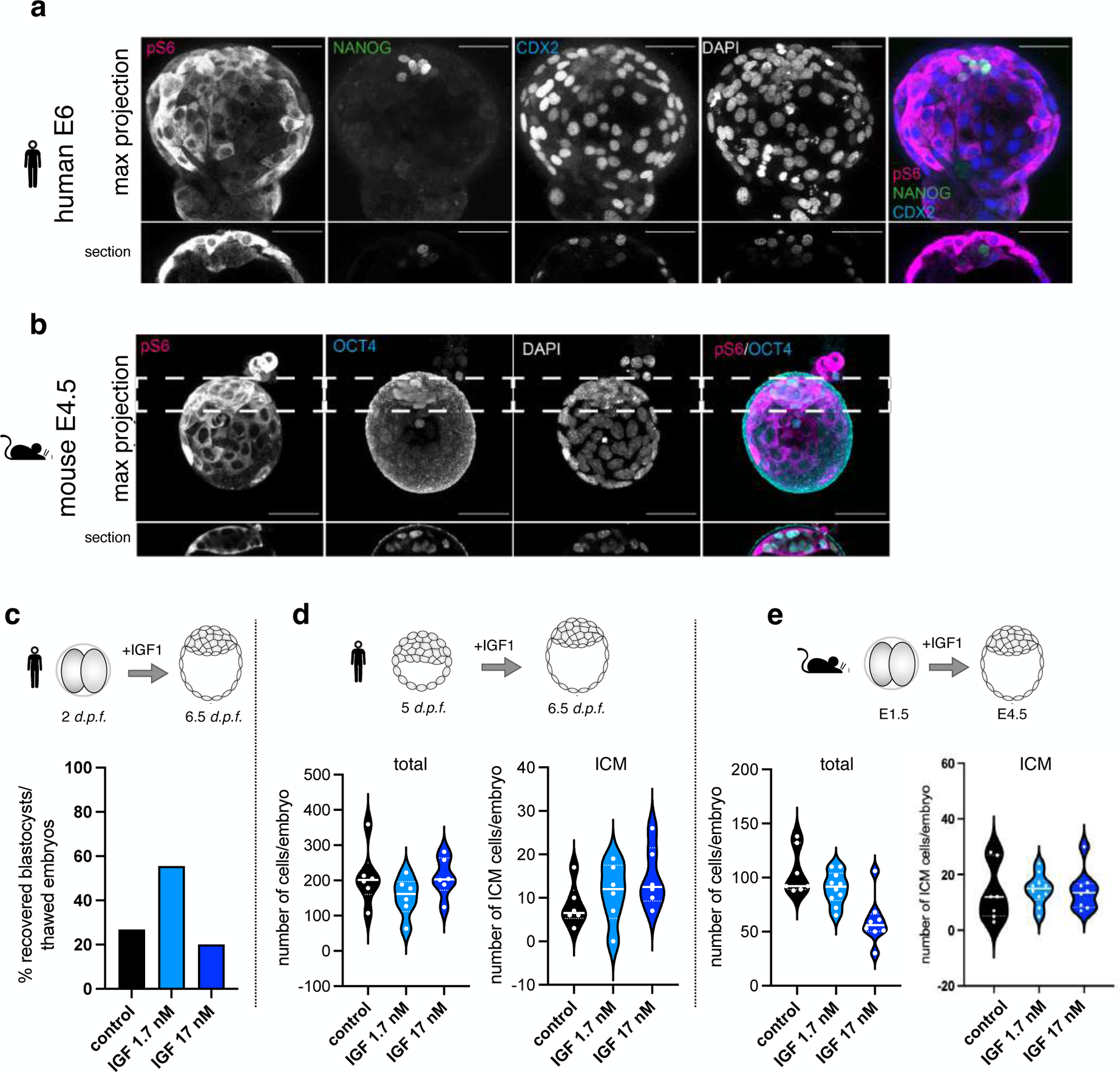
Similar pattern and sensing of mTOR pathway activity in human and mouse pre-implantation embryos. **a.** Immunofluorescence staining of a human blastocyst at E6 for the mTOR downstream target pS6, the epiblast marker NANOG, and the trophectoderm marker CDX2. Scale bars, 50 um. **b.** Immunofluorescence staining of a mouse blastocyst at E4.5 for pS6 and the ICM marker OCT4. The mouse and human embryos display a similar pattern of pS6 staining. Scale bars, 50 um. **c.** Human pre-implantation embryos cultured in 1.7 nM IGF from 2-cell until blastocyst stage yield a significantly higher percentage of blastocysts compared to control embryos. n = 14 and 18 embryos were used for 1.7 nM and 17 nM treatments, respectively. As control, data from all thawed blastocysts in the assay year were collected, n = 137. **d-e.** IGF treatment increases the number of ICM cells in human (d) and mouse (e) pre-implantation embryos.

The mTOR pathway regulates cellular growth and proliferation in response to nutrients as well as certain growth factors in the environment (*22*). The insulin growth factor (IGF) is a direct upstream regulator of the mTOR pathway and has been implicated in diapause regulation in other species (*23, 24*). IGF1 is expressed from the maternal tissue, in an estrogen-dependent manner, for paracrine regulation of the embryo, whereas the embryo expresses IGF2 for autocrine regulation (*25*). pS6 and pAKT levels are altered in uterine fibroids in response to IGF1, but not IGF2 (*26*), showing the sensitivity of mTOR downstream targets to IGF1. To test whether mTOR pathway activity is similarly sensed and adjusted in human and mouse embryos, we subjected these to two different IGF1 concentrations during the pre-implantation embryo culture (in the human either from day 2 or day 5 onwards, based on embryo availability) (Figure 1c-e). Two doses of IGF1 were used: 17 nM, the estimated concentration in the human reproductive tract (*27*), or 1.7 nM. IGF1 supplementation increased human blastocyst formation rate at 1.7 nM (58% in 1.7 nM IGF1 vs. vs 27% control media, Figure 1c), consistent with previous findings (*27*). Importantly, 1.7 nM IGF1 specifically benefitted the ICM in both human and mouse blastocysts by increasing the number of cells (Figure 1d-e, S1b). Higher IGF1 concentration did not enhance this trend, which could be explained by the increase and plateau of pS6 at 1.7 nM IGF1 in the human ICM (Figure S1c). These data suggest that the mTOR pathway is active and tightly regulated in human blastocysts in a manner that is highly similar to the mouse, supporting the possibility that mTOR inhibition may induce a diapause-like dormant state in human embryos.

### Developmental delay of human blastoids in extended pre-implantation culture

To robustly test the human diapause response in a reversible setup, we chose the recently developed blastoid model. Blastoids are embryonic stem cell-derived structures that are morphologically and transcriptionally highly similar to the human blastocyst (modeling days 5 to 7) (*28, 29*). Blastoids are generated in a relatively high-throughput manner, allowing to overcome the material and ethical limitations of human embryo research. Additionally, attachment or in vitro implantation of blastoids induce expansion and maturation of embryonic and extraembryonic cells, recapitulating these early post-implantation events and thereby allowing to test the reversibility of dormancy.

To test the effect of mTOR inhibition on the developmental timeline and morphology of human blastoids, these were first formed and then further cultured in the presence of the mTOR inhibitor RapaLink-1 (Figure 2a-b). RapaLink-1 is a third generation mTOR inhibitor, which is a rapamycin-INK128 conjugate that blocks both the allosteric and catalytic sites on mTOR and effectively blocks mTOR activity (Figure S2a). RapaLink-1 is significantly more efficient in inducing mouse diapause compared to INK128 only (*1*). RapaLink-1-treated human blastoids could be retained in their original morphology with an expanded blastocoel and an intact ICM for several days (Figure 2a-b, Figure S2b). In contrast, the majority of untreated blastoids could not be maintained beyond day 2. Over the duration of the extended culture (termed ‘pausing’ from here on), the TE of human blastoids continued to expand, whereas the ICM retained its original cell number (Figure 2b). As a result, blastoid dimensions increased compared to the untreated counterparts on day 0 (the blastoids were generated in 400 micrometer wells). The same morphological changes and TE expansion are also seen in naturally and in vitro diapaused mouse embryos (Figure 2c) and are in line with previous characterizations of embryos in diapause (*21, 30*). Thus, mTORi induces similar, diapause-like morphological changes in both human and mouse. Overall, ∼50% of human blastoids could be maintained in culture for 5 days, with a subset persisting until day 8 (Figure 2d). Paused blastoids showed markers of pluripotent and extraembryonic cells similar to untreated blastoids (Figure 2e), fulfilling a major criterion of diapause, namely stabilization of the embryo over an extended time period. Importantly, direct inhibition of protein synthesis via cycloheximide (CHX) could not replicate the effect of mTORi, indicating that mTOR function in diapause extends beyond regulation of protein synthesis (Figure S2c). Interestingly, at high doses (1000 ng/ml), CHX treatment allowed pausing of the TE at the expense of ICM (Figure S2d-e). These results corroborate the different mTOR activity levels in the TE and ICM as observed earlier (Figure 1a-b, S1).

**Figure 2.**
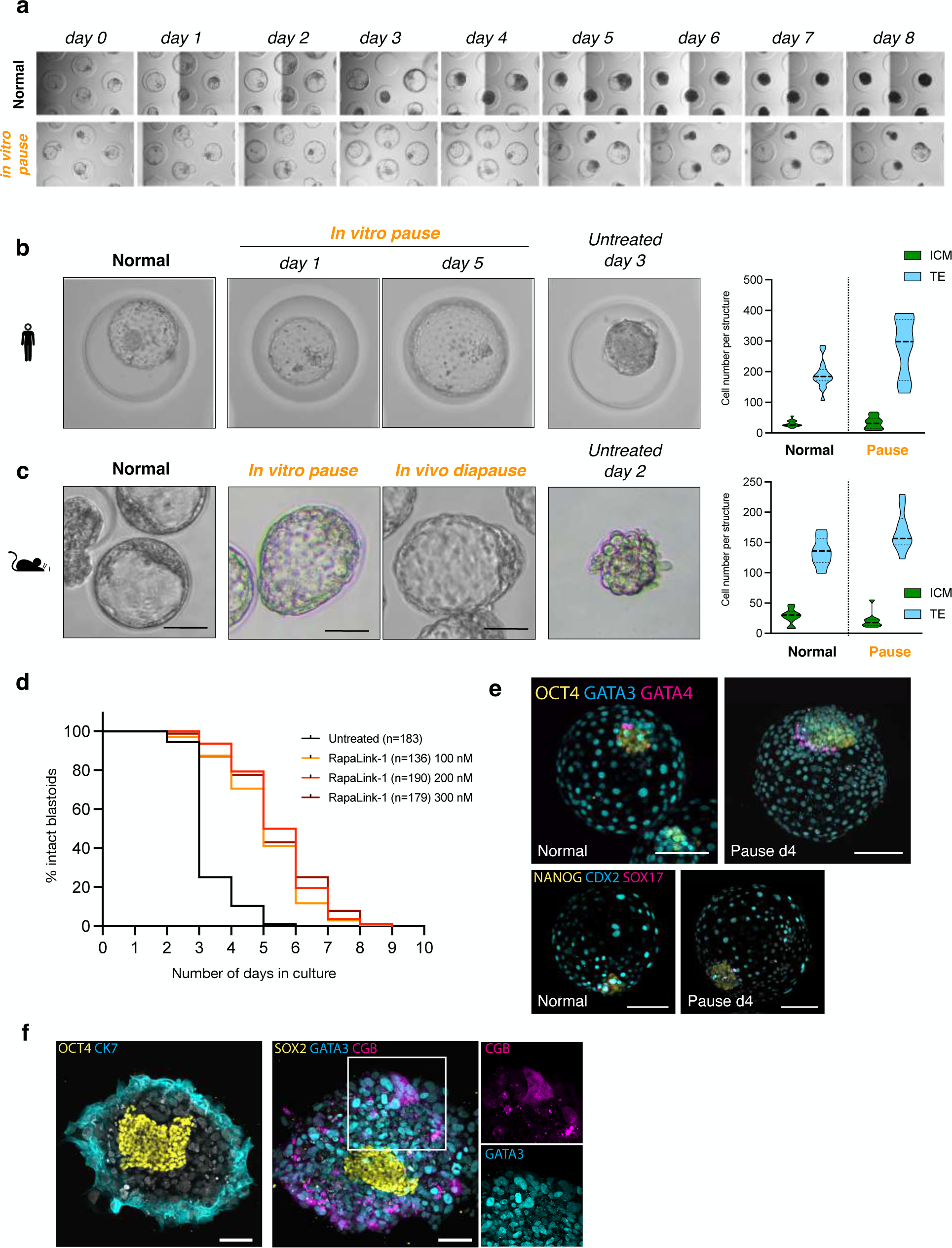
Developmental delay of human blastoids in extended pre-implantation culture. a. Bright field images of blastoids cultured in control conditions or treated with the mTOR inhibitor RapaLink-1. Indicated time points (days 1-8) denote culture time after blastoid formation (after 96h starting from ESC culture). b. Bright field images of human normal and paused blastoids, in comparison to an untreated blastoid in extended culture. Right panel shows quantification of cell number in the ICM and TE based on immunofluorescence stainings. c. Bright field images of mouse normal and paused blastocysts (in vitro and in vivo), in comparison to an untreated blastoid in extended culture. Right panel shows quantification of cell number in the ICM and TE based on immunofluorescence stainings. d. Longitudinal morphological scoring of control blastoids and blastoids treated with the mTORi RapaLink-1 at the indicated concentrations. Blastoids were not considered intact after structural collapse. n = number of treated blastoids. e. IF staining of the ICM markers OCT4 and NANOG, TE markers GATA3 and CDX2, and PrE markers GATA4 and SOX17. 16 control and 15 paused blastoids were stained. 3/15 paused and 2/16 control blastoids show linearly organized PrE layer (as shown in the top right panel). f. Reactivation and extended post-implantation culture after pausing. Blastoids were treated with mTORi for 4 days, then mTORi was withdrawn and blastoids were cultured on matrigel-coated plates (28). After four days, cells were stained for the ICM markers OCT4 and SOX2, the TE marker GATA3, and the trophoblast differentiation markers CK7 and CGB. Scale bars, 100 um.

In utero, diapaused embryos reactivate and implant upon receiving growth cues. To test whether mTORi-induced pausing is similarly reversible, we reactivated paused blastoids after 2-6 days of mTORi treatment and further cultured them on matrigel-coated plates (Figure 2f, S3). The efficiency of attachment was 100% in all conditions of reactivation. ICM, TE and primitive endoderm (PrE) derivatives were detected after 2-4 days of post-implantation culture (Figure 2f, S3), including further differentiating TE derivatives that express CK7 or CGB (Figure 2f). We noted a skewing towards PrE fate after 6 days of pausing, although this may depend on the specific reactivation conditions (Figure S3). These results show that the pausing is reversible and the capacity to reactivate and further differentiate is retained in human blastoids, thus fulfilling a further criterion of diapause.

### Human naïve and naïve-like PSCs can adopt a dormant state

We have previously shown that mouse naïve pluripotent stem cells (PSCs) under mTORi can faithfully model the diapaused epiblast, including high transcriptional similarity (*1*). Therefore, to further corroborate and dissect the human diapause response, we investigated the possibility of human PSCs to adopt a dormant state (Figure 3).

**Figure 3.**
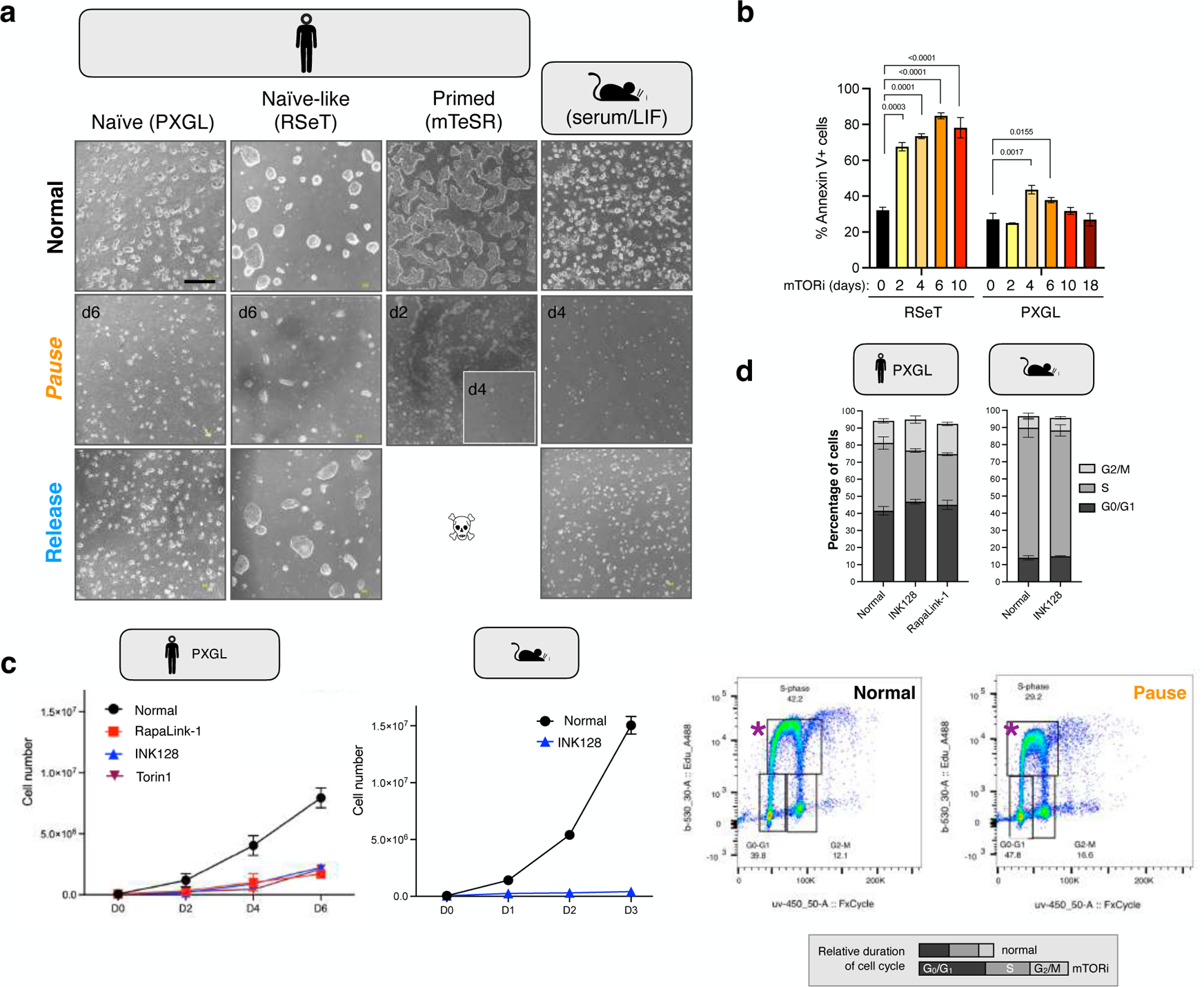
Human pluripotent cells can be reversibly paused with differing efficiencies via inhibition of mTOR activity. **a.** Representative bright field images of naïve, naïve-like, and primed pluripotent human cells before, during, and after mTORi treatment as compared to the mouse. Scale bar, 500 um. **b.** Percentage of Annexin V-positive apoptotic cells during the course of mTORi treatment in RSeT and PXGL culture. **c.** Proliferation curves of human ESCs in PXGL culture treated with the catalytic mTOR inhibitors RapaLink-1 and INK128. Proliferation rate of mouse ESCs cultured in INK128 are shown in comparison. **d.** Cell cycle distribution of mTORi-treated human and mouse cells, as determined by EdU incorporation and DNA content. Differences are non-significant per two-way ANOVA. Schematic at the bottom shows the relative durations of the cell cycle and each phase, based on measurements of proliferation and EdU integration (Figure 3c-d).

Both mouse and human PSCs can be captured in vitro with molecular and epigenetic signatures and morphological properties of either naïve or primed pluripotency that respectively reflect the blastocyst (*31–41*) and the early post-implantation epiblast (*42*). Several groups have established culture conditions to maintain human PSCs in a naïve state (*31–41*). Since diapause in most mammals occurs before implantation, naïve PSCs may be more amenable to be put in a diapause-like state. To comparatively test the capacity of PSCs across the pluripotency spectrum to enter dormancy, we used human PSCs cultured in naïve (PXGL), naïve-like (RSeT) or primed (mTeSR) conditions. Mouse ESCs in serum/LIF, which we previously showed to robustly transition into a diapause-like state in response to mTORi, were used as control. As a first measure, we observed morphology, proliferation rate, and cell death during the potential cellular transition from a proliferative state to dormancy. As expected, mouse ESCs successfully slowed their proliferation while maintaining colony morphology, and this effect was reversible (Figure 3a).

Strikingly, human naïve PSCs cultured in PXGL conditions showed a similar response to mouse ESCs and robustly adopted a less proliferative pluripotent state that was reversible. Naïve PSCs could be maintained under mTORi for 18 days (maximum testing period), released and iteratively repaused without compromising colony morphology (Figure S4a). Cells cultured in RSeT could also be reversibly paused, albeit at much lower efficiency. While PXGL cells showed a more homogeneous response to mTORi treatment, with an initial increase in apoptosis that normalized over time, RSeT cells showed persistently high levels of apoptosis (Figure 3b). Yet, the remaining RSeT colonies retained pluripotent morphology, did not accumulate DNA damage, and could also revert to proliferation without differentiation (Figure 3a, S4b). Furthermore, paused RSeT colonies showed stronger dormancy and effectively abolished proliferation while still maintaining pluripotency (Figure S5). In contrast to PXGL and RSeT cells, primed PSCs in mTeSR culture did not successfully transition into dormancy and died by day 4 (Figure 3a). These results reveal that human cells can undergo reversible dormancy in response to mTORi in naïve and naïve-like states, with naïve cells showing higher efficiency and naïve-like cells entering deeper dormancy.

Mouse and human PSCs were so far paused using the mTOR inhibitor INK128 as this was previously shown to effectively instate a diapause-like state in mouse ESCs (*1*). INK128 efficiently inhibited the phosphorylation of mTOR downstream targets pS6 and Akt in mouse and human cells, albeit pS6 more then pAkt (Figure S6). Treatment of PXGL cells with two other mTOR inhibitors, RapaLink-1 and Torin1, exactly reproduced the slowed proliferation under INK128 (Figure 3c). Interestingly, proliferation of human PSCs was less suppressed than mouse ESCs using the same inhibitor (INK128), which may be attributed to a longer growth (G_1_) phase (Figure 3c-d). Of note, even though paused hPSCs in PXGL showed an overall similar cell cycle distribution, the nucleotide analog EdU was integrated at lower levels in paused cells, indicating slower progression through the cell cycle (Figure 3d, indicated by asterisks) Overall, the cell cycle and proliferation data point to a prolonged cell cycle in PXGL cells, in which all phases take longer to complete, with G_0_/G_1_ and G_2_/M phases prolonging proportionally more than the S phase compared to normal cells (depicted in Figure 3d, lower panel). In contrast, paused RSeT cells did not show the mitosis marker H3S10p (Figure S5).

### Reversibility of pausing at the molecular level

Our results so far indicate that human PSCs as well as blastoids have the capacity to pause and that pausing is functionally reversible. We next aimed to probe the reversibility of pausing at the molecular level, identify the involved pathways, and more thoroughly compare PXGL and RseT conditions. For this, we collected normal, stably paused, and released cells and performed quantitative label-free proteomics mass spectrometry (Figure 4a, Table S1). Principle component analysis (PCA) showed a clearly distinct proteomics state in paused PSCs, which was completely reversible in RseT cells and showed more heterogeneity in PXGL cells (Figure 4b). 321 and 392 proteins were significantly differentially expressed (DE, adjusted p-value <0.05 and log_2_FC >1) in PXGL and RseT conditions, respectively (Table S2). Both at the pathway and individual protein levels, pause-induced changes reverted back to original after release of cells into proliferation (Figure 4c-d, Table S3). Again, RSeT cells showed almost completely restored original expression levels and patterns, while PXGL response was more variable. Protein synthesis, which is a major downstream target of mTOR, was consistently downregulated in both PXGL and RSeT cells, along with other anabolic activities such as transcription, splicing, and RNA/protein export. Upregulated pathways differed to some extent under these conditions (Figure 4c). Overall, these results reiterate the reversibility of pausing in human PSCs with differing efficiencies at the functional and molecular levels between naïve and naïve-like conditions.

**Figure 4.**
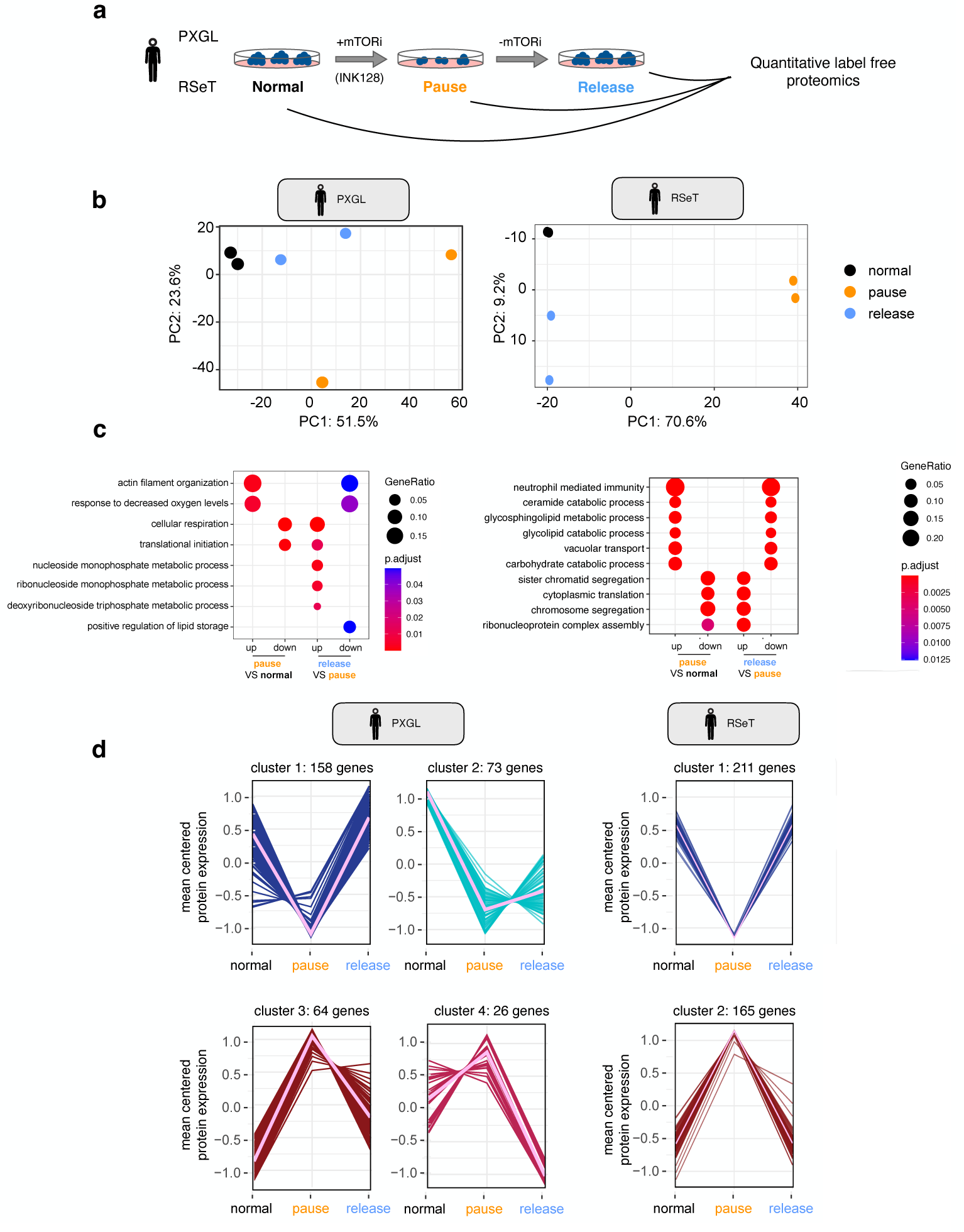
Reversibility of human PSC pausing at the molecular level. **a.** Schematics of the experiment. Normal, paused, and released human PSCs in PXGL and RSeT conditions were collected and subjected to quantitative label-free mass spectrometry. **b.** Principal component analysis of top 500 variable proteins in the indicated conditions in PXGL and RSeT culture. **c.** Gene ontology analysis of differentially expressed proteins in paused and released PXGL and RSeT cells. Unique terms are shown to reduce redundancy. Complete lists are provided in Table S3. **d.** K-means clustering of downregulated (left) or upregulated (right) genes during pausing shows reversibility of expression pattern. Scaled log2(LFQ+1.1) values are shown. LFQ, label-free quantification.

### Species-specific tempo and metabolism of developmental pausing

To get at a potentially conserved diapause response in mammals, we next compared the temporal dynamics and pathway usage in mouse and human naïve PSCs in their cellular transition into dormancy. For this we collected matching time points up to 10 days in mouse and human and performed proteomics (Figure 5a).

**Figure 5.**
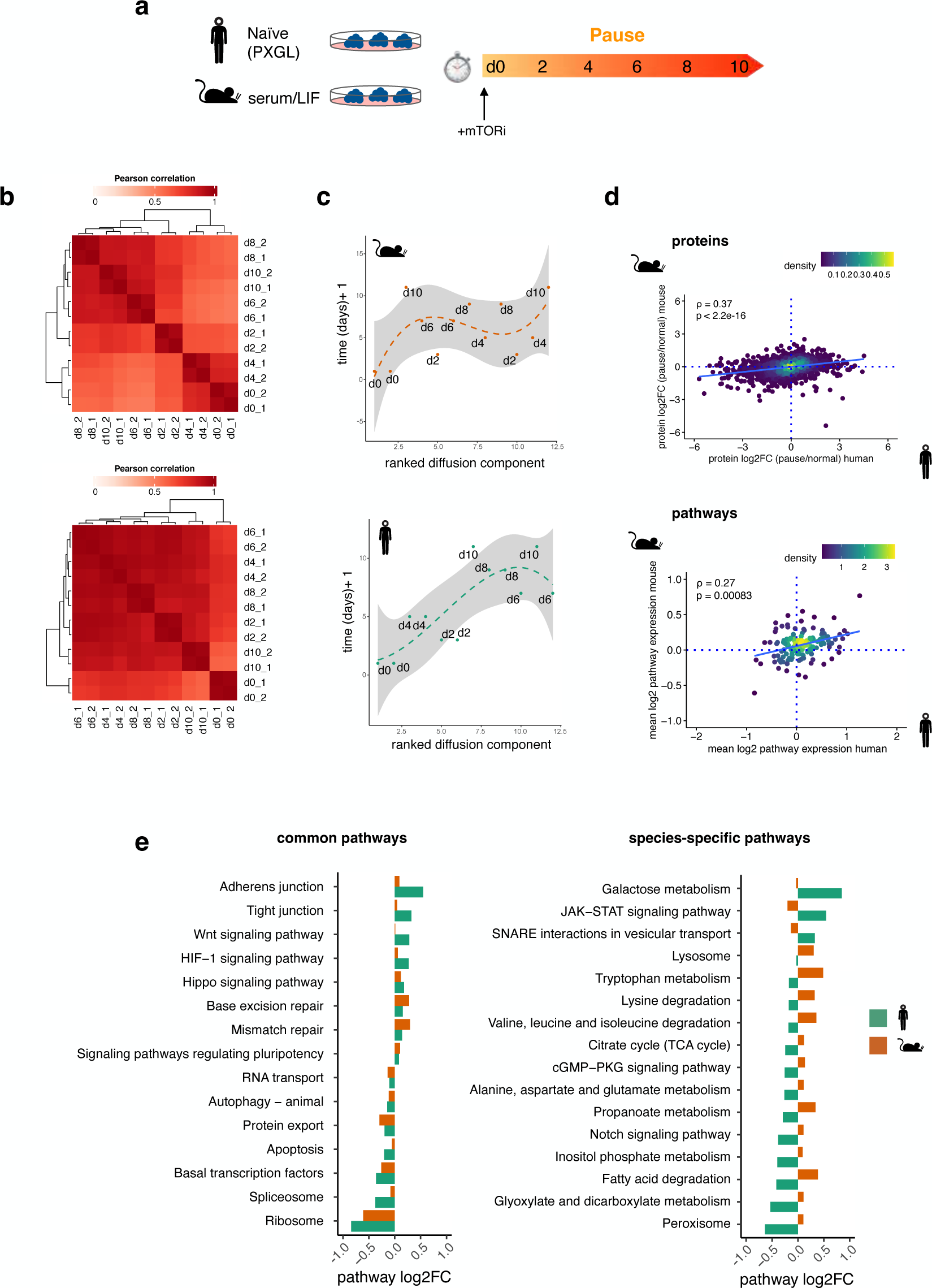
Distinct temporal dynamics and metabolic pathway expression in human and mouse pausing. **a.** Schematics of the experiment. Human ESCs in PXGL culture and mouse ESCs in serum/LIF culture were treated with mTORi for 10 days and cells were collected at the indicated time points for low-input mass spectrometry. To investigate reversibility, cells were released from pausing and collected 8 days after. Mouse and human cells were cultured in parallel using the same batch of the inhibitor. **b.** Correlation matrices of human (PXGL) and mouse ESCs. Two replicates (dx_1 or _2) were collected on the indicated days for proteomics analyses. **c.** Pseudotime analyses of human or mouse ESCs undergoing pausing based on ranked diffusion components. Dashed line and grey shading show median and confidence interval, respectively. Each dot shows a replicate collected on the indicated day of mTOR treatment. **d.** Density plots showing the levels of protein (top) or pathway (bottom) expression in human and mouse cells in pausing relative to normal culture. Proteomic and pathway alterations in response to pausing show a statistically significant correlation in both species. r, Spearman’s Rho correlation coefficient. **e.** Common and species-specific pathways in human and mouse paused ESCs. Basal transcription and translation pathways are down-regulated; while DNA repair proteins are upregulated in both species. Metabolic profiles are strikingly species-specific.

The developmental clock of mice and humans is inherently different and we thus wondered whether diapause is established at a similar or distinct pace in these species. Correlation matrices of longitudinal samples showed different ‘break points’ in human and mouse cells, after which the paused proteomic signature was stabilized (human = d6, mouse = d2, Figure 5b). To corroborate this observation, we further used the time-course proteomics data to construct pseudotime trajectories using ranked diffusion components. This approach also pointed to d6 in human and d2 in mouse as the observable time points, after which the pseudotime no longer consistently progressed, and thus stabilized (Figure 5c). These results strongly suggest a species-specific tempo in PSCs of transition into dormancy.

Once stabilized, human and mouse pause proteomes correlated at the protein and pathway levels (Figure 5d). Here we chose the time points after the stabilization of pseudotime, i.e. day 8 in human and day 4 in mouse, with the aim of identifying pathways important for the *maintenance* of pausing. In general, RNA and protein anabolic processes were downregulated in both species (e.g. ribosome, basal transcription, RNA and protein export), whereas cellular adhesion and DNA repair-related terms were commonly upregulated (Figure 5e). Despite an overall correlation, we also identified several species-specific pathways. These mostly comprised of metabolic pathways, with lipid and amino acid metabolism specifically upregulated in the mouse; and sugar metabolism specifically upregulated in the human (Figure 5e). Only a few signaling pathways were discordantly expressed in the two species (Figure 5e). These differences might however also stem from different culture conditions, e.g. PXGL media contains a WNT inhibitor, unlike mouse serum/LIF media. Taken together, the human and mouse response to mTORi-induced pausing builds on a common non-anabolic profile, but may require species-specific metabolic activities.

Finally, we sought to validate the species-specific metabolic requirements, as predicted by the proteome analyses of paused human PSCs, using the blastoid model. Proteomic analysis pointed to fatty acids as a mouse-specific energy source utilized to sustain pausing (Figure 5e, S7a). We and others have recently shown that, indeed, fatty acids derived from lipid droplets are actively used during mouse diapause (*20, 43*), and that enhancing fatty acid oxidation by L-carnitine supplementation significantly prolongs in vitro pausing duration of mouse blastocysts (*20*). Consistent with species-specific energy sources, L-carnitine supplementation of human blastoids during pausing did not alter the culture duration (Figure S7b). The neutrality to carnitine supplementation suggests a non-lipid-based metabolism in paused human blastoids. Understanding and modulating the metabolic profile of human diapause may thus further enhance the pausing efficiency and duration of human blastoids, and potentially, blastocysts.

## DISCUSSION

Human embryos are notoriously variable in their capacity to progress through early development, with an estimated 1/3 of pregnancies failing in the first trimester due to genetic and non-genetic abnormalities (*44, 45*). In this context, our results have several important implications for technological advance as well as fundamental understanding of human early embryogenesis: 1-Stimulation of mTOR/PI3K pathway activity through supplementation with IGF1 may promote cell proliferation and boost blastocyst formation rates, supporting earlier findings (*46–48*); 2-Inhibition of the mTOR pathway at the blastocyst stage might provide a valuable extended time window for characterization and scoring of embryos or synchronization to the mother’s hormonal cycle; 3-paused hPSCs and blastoids can be used to understand factors regulating the prolonged maintenance of pluripotency (similar to the role of LIF in mouse pluripotency (*30*)).

Once established and stabilized, paused hPSCs and blastoids easily reactivate upon withdrawal of mTORi. However, the starting population of naïve and naïve-like hPSCs show heterogeneity in terms of the capacity to pause. The underlying reasons for this heterogeneity remain to be explored. We speculate that transcriptional status or the cell cycle phase of the cells at the time of entry to pausing could be contributing factors (*49*). Analysis of transcriptome profiles at the single cell level and/or lineage tracing experiments may help identifying the causes of heterogeneity. Since the cells are isogenic, the contribution of different genetic factors cannot be isolated in this setting. We note that human PSCs have an increased tendency for apoptosis, and unlike mouse PSCs, require periodic use of ROCK inhibitor for stabilization, particularly at the time of passaging. Thus, maintaining hPSCs under mTORi is more challenging as compared to mouse cells. Among the conditions tested in this manuscript, PXGL media offers a more stable platform for further investigations on human embryonic dormancy. Yet, it is advisable to compare different pluripotent states as well as use blastoids for functional testing of critical regulators of dormancy.

In the recent years, regulated translation emerged as an important new player connecting stem cell activity to the microenvironment (*50–53*). Downregulation of translation is the most prominent signature of hPSC pausing and correlates with dormancy. However, in mouse, neither embryos nor cells can be paused by inhibition of translation alone (*1*). The same is true for human blastoids, which interestingly lose the epiblast upon prolonged inhibition of protein synthesis. These results strongly suggest that the mTOR pathway activity is stringently controlled in a tissue-specific manner. mTORi-based rewiring of metabolism appears to be a necessary component for maintenance of the dormant state. Upstream of mTOR, selective depletion of nutrients or growth factors such as IGF1 could trigger mTOR inhibition and developmental pausing in humans. As the expression of IGF1 is estrogen-dependent, and estrogen deprivation is the trigger for the non-receptivity of uterus, which induces diapause in the mouse; IGF1 depletion may be a main upstream regulator of diapause. Although IGF also triggers signaling pathways other than mTOR/PI3K, here we show that mTOR pathway senses IGF1 levels.

## ACKNOWLEDGMENTS

We thank members of the Bulut-Karslioglu Lab, Ludovic Vallier, Michelle Percharde and Helene Kretzmer for critical feedback, MPIMG scientific facilities for excellent service, Jennifer Shay and Cordula Mancini for assistance, Beata Lukaszewska-McGreal for proteome sample preparation. This project was supported by the German Academic Exchange Service (DAAD) PhD Fellowship to DPI (91730547), the Swiss National Science Foundation Early Postdoc.Mobility fellowship (P2EZP3_195682) to VAvdW, the Wellcome HDBI initiative (UK Human Developmental Biology Initiative 360 G-Wellcome-215116_Z_18_Z) to TR, the European Research Council (ERC) under the European Union’s Horizon 2020 research and innovation programme (ERC-Co grant agreement no.101002317 ‘BLASTOID: a discovery platform for early human embryogenesis’) to NR, the Austrian Science Fund (FWF) through a Lise Meitner Programme (project no. M3131-B) to HHK, the Max Planck Society (EGS and ABK) and the Sofja Kovalevskaja Award (Humboldt Foundation) to ABK. Work in the laboratory of KKN was supported by the Francis Crick Institute, which receives its core funding from Cancer Research UK (FC001120), the UK Medical Research Council (FC001120), and the Wellcome Trust (FC001120). For the purpose of Open Access, the author has applied a CC BY public copyright licence to any Author Accepted Manuscript version arising from this submission.

## AUTHOR CONTRIBUTIONS

ABK and DPI conceived the project. DPI established the human PSC pausing models and performed all stem cell experiments. VAvdW performed computational analysis and mouse embryo stainings. HHK performed blastoid experiments under the guidance of NR. ID and EGS shared resources. AM, TR, CSS, and SEW performed human embryo experiments under the supervision of KKN. KE, PS, and LC coordinated the donation of embryos. ABK supervised the project. DPI, VAvdW, NR, and ABK wrote the manuscript with feedback from all authors.

## SUPPLEMENTARY FIGURES

**Figure S1.**
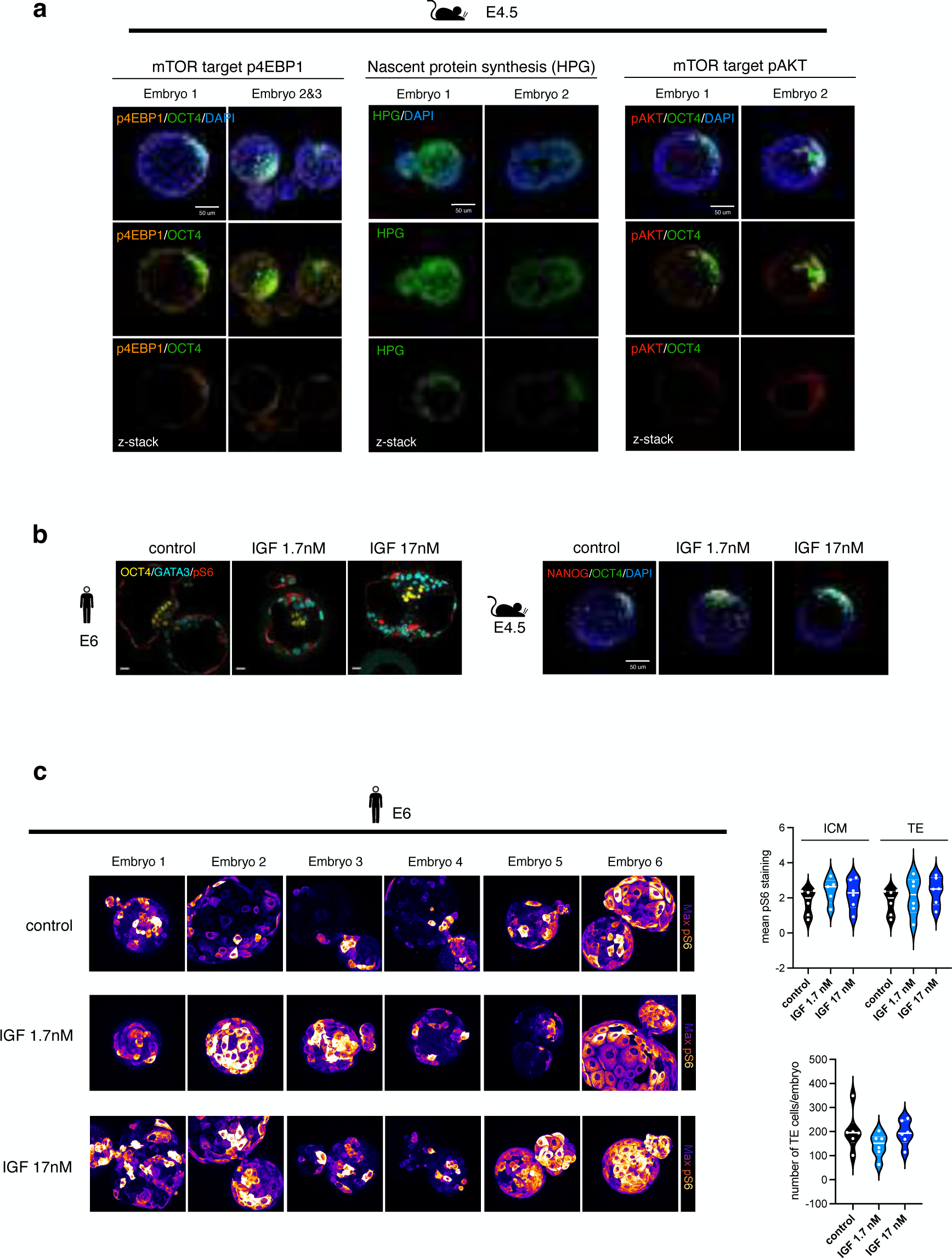
Characterization of mTOR pathway in mouse and human blastocysts. **a.** Immunofluorescence stainings of the mTOR downstream targets pAKT and p4EBP1 and nascent protein synthesis in mouse E4.5 embryos. Embryos were cultured with the amino acid analog HPG was used to mark nascent proteins. All three stains show heterogeneity in pathway activity between ICM and TE and within TE. **b.** IF stainings showing ICM and TE in human and mouse blastocysts in control or IGF treatment conditions. These data support Figure 1d-e. **c.** IF staining for pS6 in human pre-implantation embryos in control condition and upon treatment with IGF1. Fluorescence maximum intensity projections are shown. Right panels show quantification of pS6 intensities and number of TE cells in each condition. ICM reaches maximum pS6 intensity in 1.7 nM IGF1 treatment. Solid lines indicate median and dashed lines mark interquartile range.

**Figure S2.**
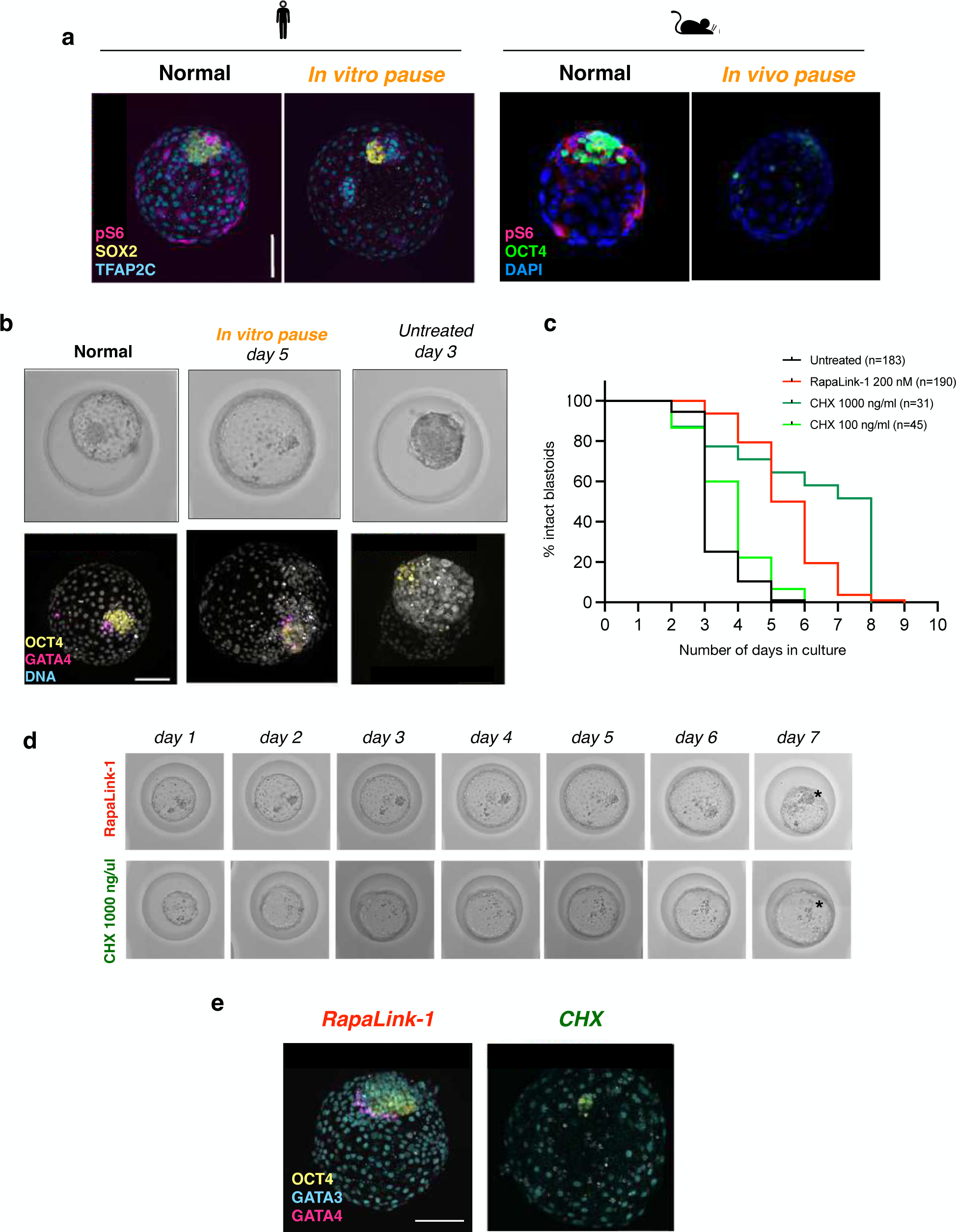
Further characterization of blastoid pausing. **a.** Immunofluorescence stainings for pS6 in human normal and paused blastoids (left) and mouse normal and in vivo diapaused blastocyst. Note that pS6 staining is completely absent in the paused states. **b.** Immunofluorescence stainings of the epiblast (OCT4) and PrE markers (GATA4) in normal, paused, and untreated blastoids. **c.** Longitudinal morphological scoring of control blastoids and blastoids treated with RapaLink-1 or CHX (protein synthesis inhibitor). Blastoids were not considered intact after structural collapse. n = number of treated blastoids. **d.** Bright field images of selected blastoids cultures in the indicated conditions. Note the loss of ICM (indicated with asterisk) in CHX-treated blastoids. **e.** Immunofluorescence staining corroborating the premature loss of OCT4+ epiblast cells in CHX-treated blastoids on day 4.

**Figure S3.**
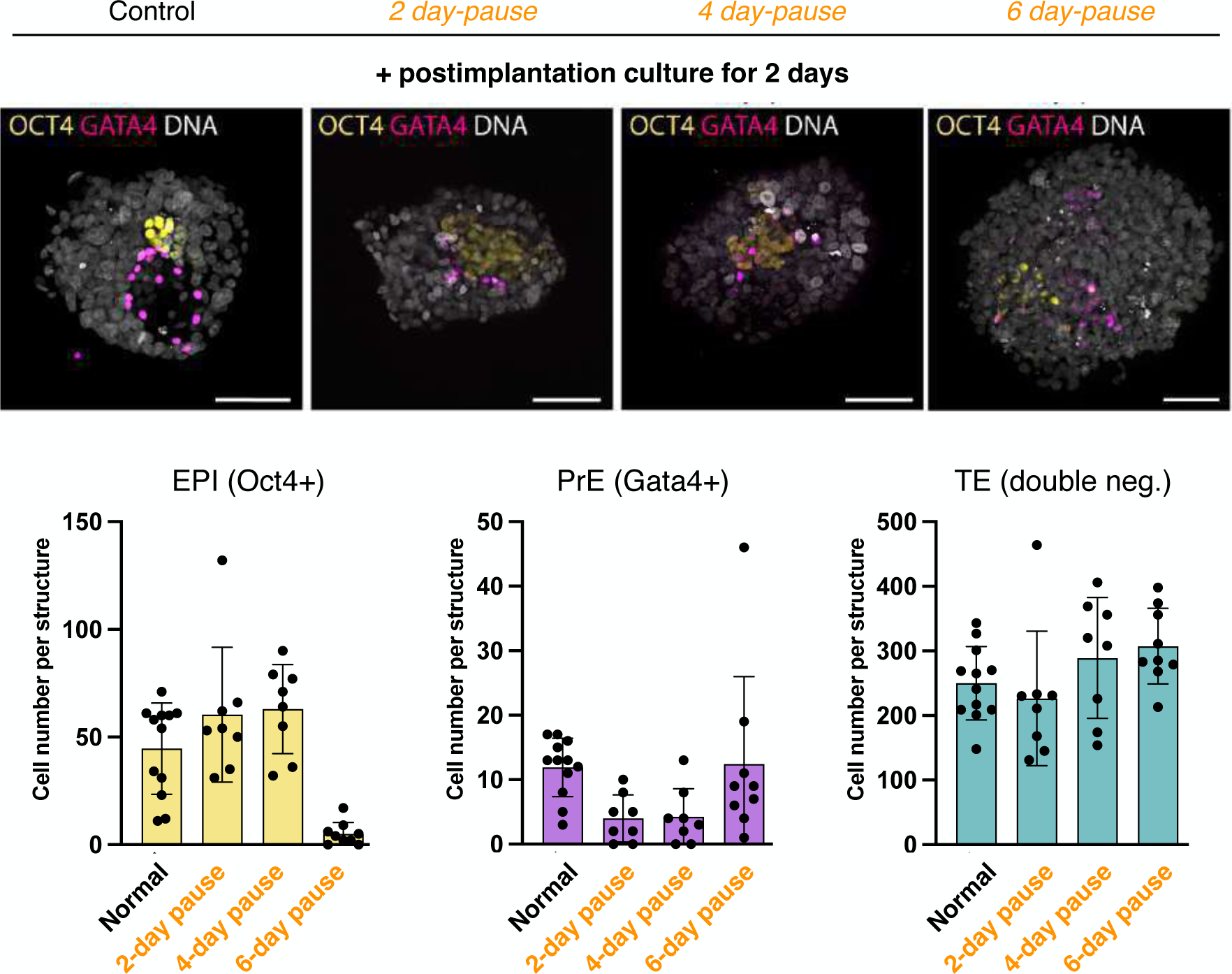
Further characterization of blastoid reactivation. **a.** Reactivation and extended post-implantation culture after pausing. Blastoids were treated with mTORi for 2-6 days, then mTORi was withdrawn and blastoids were cultured on matrigel-coated plates(*28*). After two days, cells were stained for the ICM marker OCT4 and the PrE marker GATA4. Number of cells in each lineage was quantified (TE was defined as double negative). Scale bars, 100 mm.

**Figure S4.**
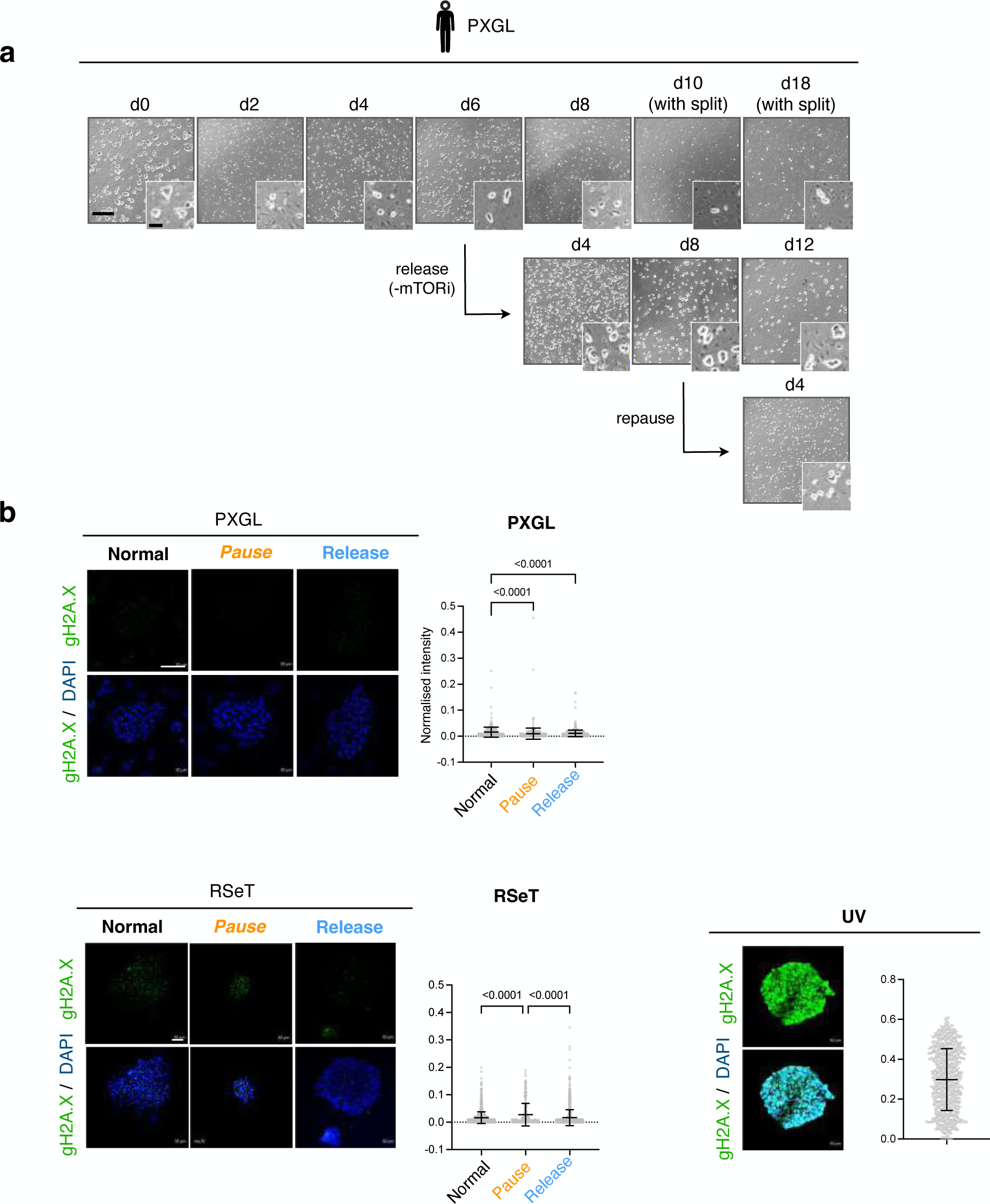
Further characterization of paused hPSCs in PXGL and RSeT culture. **a.** Bright field images of paused, released, and re-paused human ESCs in PXGL culture. Cells need to be split during prolonged pausing to overcome MEF depletion and differentiation. Scale bars: main-500 mm, inset-100 mm. **b.** Levels of the DNA damage marker gH2A.X in the corresponding conditions. As positive control, cells were irradiated with UV at 4000mJ/sq cm for 10 seconds and fixed 6 hours later. Paused and released hiPSCs do not show an increase in DNA damage or apoptosis. Insets show higher magnification of the indicated regions. Right panels show single-cell quantifications of gH2A.X intensity normalized to the cellular area. Mean and standard deviation are shown. n = number of cells. Scale bars, 50 mm. Statistical test performed is one-way ANOVA with Tukey’s multiple testing correction).

**Figure S5.**
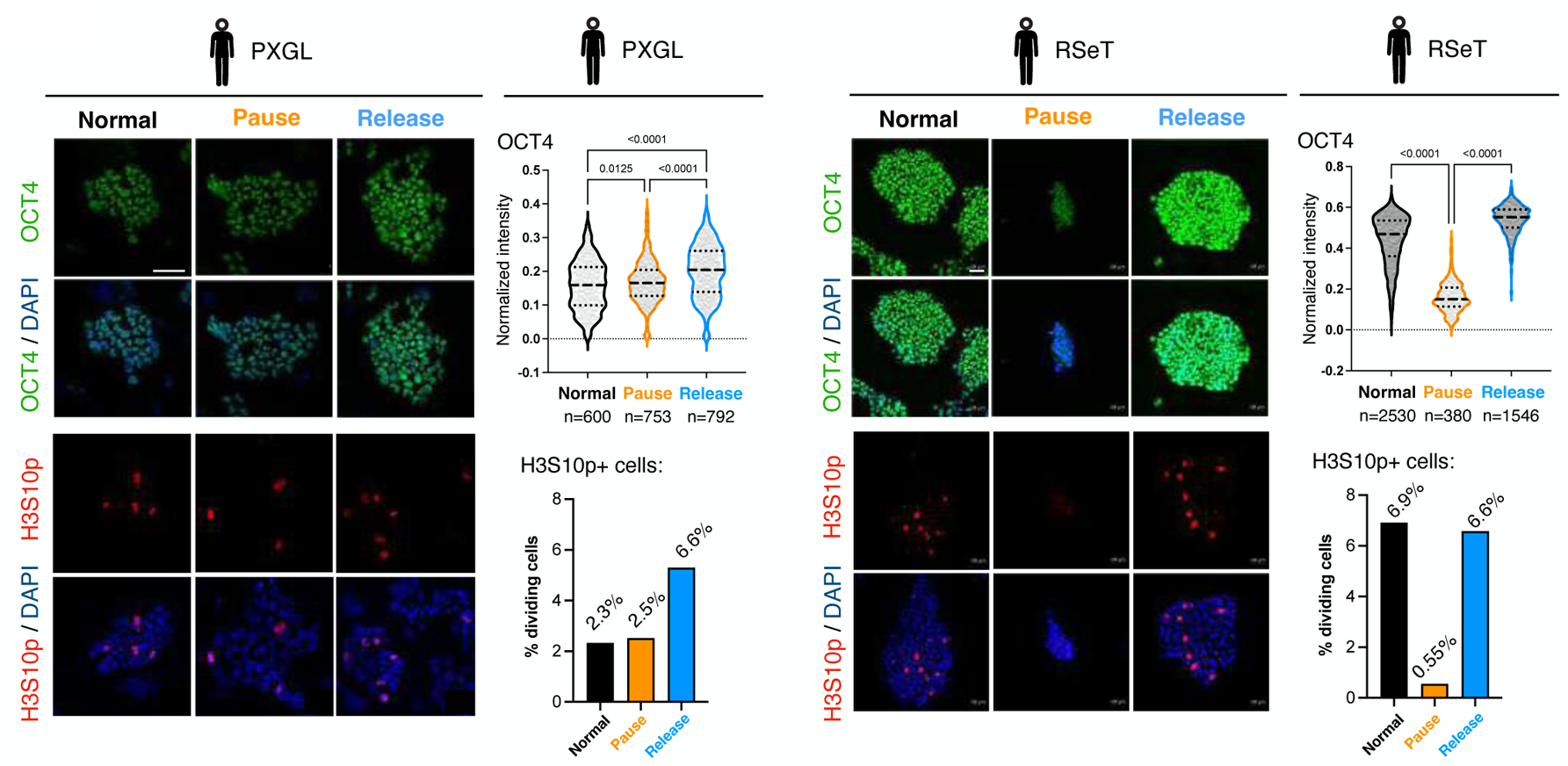
Paused hPSCs retain pluripotency and the capacity to proliferate. Levels of the pluripotency-associated gene OCT4 and the cell division marker H3S10p in normal, paused (d6), and released human cells in PXGL and RSeT culture. Right panels show single-cell quantifications all taken images. n indicates the number of analyzed cells. Scale bars, 50 mm. Statistical test performed is one-way ANOVA with Tukey’s multiple testing correction.

**Figure S6.**
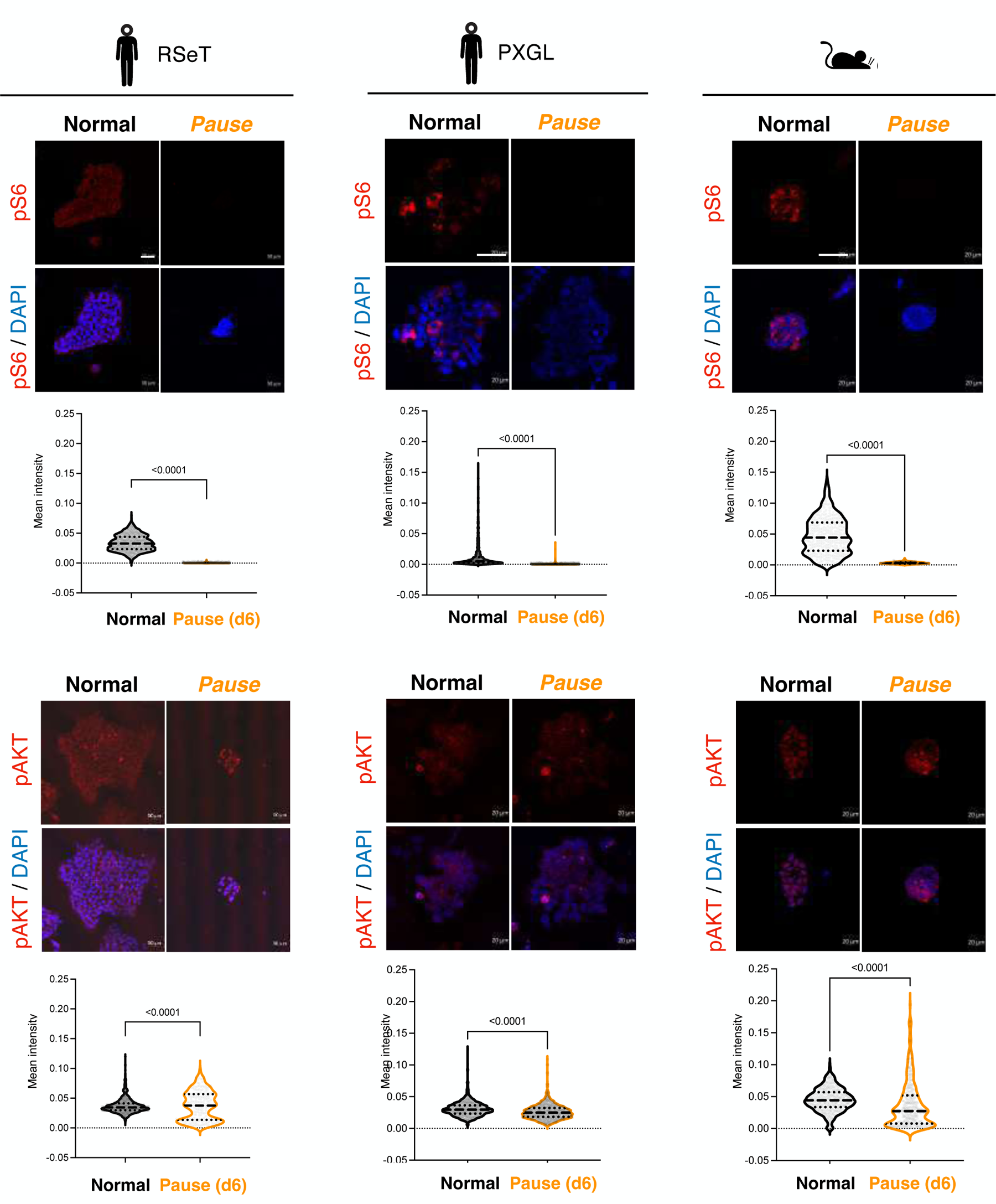
Levels of mTOR downstream targets in paused mouse and human cells. Immunofluorescence stainings for the mTOR downstream targets pS6 and pAKT. Bottom panels show area-normalized single-cell quantifications of staining intensities. Dashed line: median, dotted lines: quartiles. Scale bars, 50 mm. Statistical test performed is a two-tailed Kolmogorov-Smirnov t-test.

**Figure S7.**
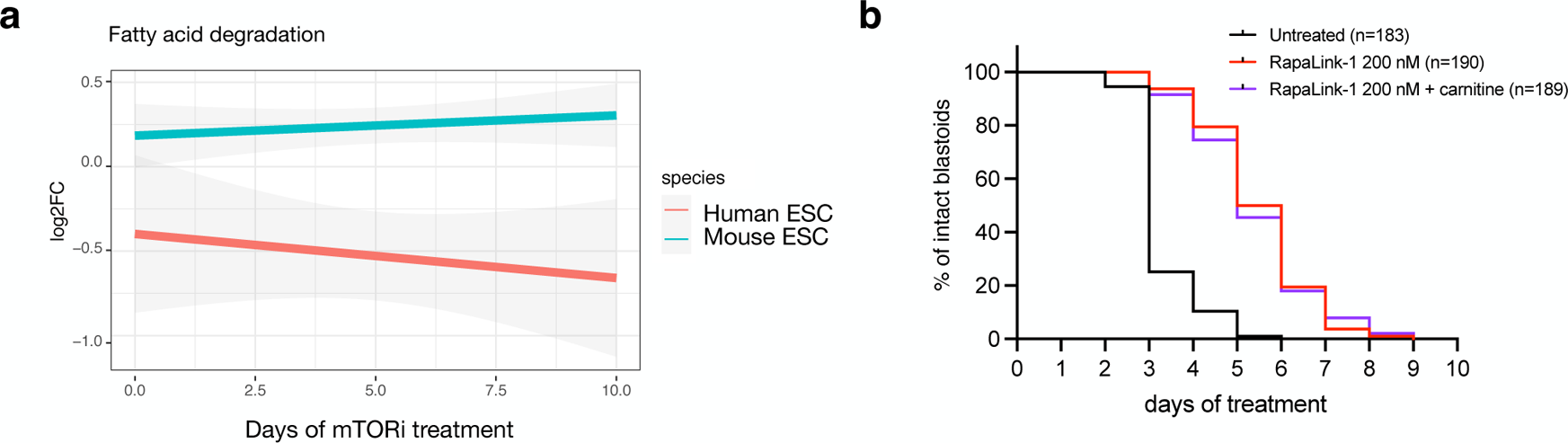
Human blastoid maintenance in pausing does not require fatty acid oxidation. **a.** Temporal expression of the fatty acid degradation pathway during mTORi treatment in mouse and human ESCs. Plot shows mean log2FC of proteins in the pathway (per KEGG annotation) on dx/d0, together with confidence interval. **b.** Culture duration analyses of paused blastoids with or without carnitine supplementation. Enhancing fatty acid degradation does not prolong pausing of human blastoids, unlike mouse blastocysts. n = number of treated blastoids.

## METHODS

### Ethics statement

Human embryos were donated to the research project by informed consent under UK Human Fertilization and Embryo Authority (HFEA) License number R0162. Approval was also obtained from the Health Research Authority’s Cambridge Central Research Ethics Committee, IRAS project ID 272218 (Cambridge Central reference number 19/EE/0297). The approval process entailed independent peer review along with approval from both the HFEA Executive Licensing Panel and the Executive Committees. Our research is compliant with the HFEA Code of Practice and has undergone independent HFEA inspections since the license was granted. Patient consent was obtained from Bourn Hall Clinic. Informed consent was obtained from all couples that donated surplus embryos following IVF treatment. Before giving consent, donors were provided with all of the necessary information about the research project, an opportunity to receive counselling, and details of the conditions that apply within the license and the HFEA Code of Practice. Specifically, patients signed a consent form authorizing the use of their embryos for research including stem cell derivation and for the results of these studies to be published in scientific journals. No financial inducements were offered for donation. Patient information sheets and the consent documents provided to patients are publicly available (https://www.crick.ac.uk/research/a-z-researchers/researchers-k-o/kathy-niakan/hfea-licence/). Embryos were donated, cryopreserved, and transferred to the Francis Crick Institute where they were thawed and used in the research project.

### Human embryo culture

Vitrified embryos frozen in straws were thawed by quickly transferring the contents of the straw from liquid nitrogen directly into thaw solution (Irvine Scientific Vitrification Thaw Kit) and thawed per manufacturer’s instructions. Embryos frozen in cryopets were thawed for 3 seconds in a 37 °C water bath before transferring into thaw solution (Irvine Scientific Vitrification Thaw Kit). Embryos frozen in glass ampoules were thawed completely in a 37 °C water bath after the top of the vial was removed under liquid nitrogen. The contents were emptied onto a petri dish and the embryo transferred through a gradient of sucrose solutions (Quinn’s Advantage Thaw Kit, Origio) per manufacturer’s instructions. Embryos were routinely cultured in Global Media supplemented with 5 mg/mL LifeGlobal Human Protein Supplement (both LifeGlobal) pre-equilibrated overnight in an incubator at 37 °C and 5% CO_2_. These conditions were supplemented with IGF1 (291-G1/ CF, R&D) at a final concentration of 1.7 or 17 nM.

### Human ESC and blastoid experiments

As required by the German Stem Cell Act, all experiments involving hESCs and/or hESC derived blastoids were approved by Robert Koch Institute, Berlin, and the Commission for Science Ethics of the Austrian Academy of Sciences. The Wicell line H9 was used under the agreement 20-WO-341 for a research program entitled ‘Modeling early human development: Establishing a stem cell based 3D in vitro model of human blastocyst (blastoids)’. This work did not exceed a developmental stage normally associated with 14 consecutive days in culture after fertilization even though this is not forbidden by the ISSCR Guidelines as far as embryo models are concerned. All experiments complied with all relevant guidelines and regulations, including the 2021 ISSCR guidelines that forbid the transfer of human blastoids into any uterus (*1*).

### Mouse experiments

All animal experiments were performed according to local animal welfare laws and approved by local authorities (covered by LaGeSo licenses ZH120, G0284/18, and G021/19 and UK Home Office project license (70/8560)) within the conditions of the Animal (Scientific Procedures) Act 1986. Mice were housed in individually ventilated cages and fed ad libitum.

### Stem cell lines and culture conditions

#### Primed human PSC cell culture

Wild-type Zip13k2 female hiPSCs were cultured without feeders on Matrigel-coated plates (Corning, 354277) in mTeSR media supplemented with 10 mM ROCK inhibitor (ROCKi) (Tocris, 1254) on the day of seeding. ROCKi was withdrawn the day after thawing, the media was changed every day, and cells were passaged every four days. At each passage, cells were dissociated as clumps using EDTA and re-plated with 10 mM ROCKi in a split ratio of 1:5. Cells were maintained at 37°C in 20% O_2_ and 5% CO_2_ incubator.

#### Naïve-like human PSC cell culture (RSeT)

To obtain naïve-like cells, primed hiPSCs were plated at medium density (1:3-1:4 of a 75% confluent 10 cm plate) on Matrigel-coated dishes (Corning, 354277) and grown in mTeSR media for 24 hours at 37°C in 20% O_2_, 5% CO_2_. After 24 hours, mTeSR media was replaced with RSeT^TM^ Feeder-Free Medium (Stem Cell Technologies, 05975), and the cells were cultured at 37°C in 5% O_2_, 5% CO_2_ (hypoxic condition). The cells need to undergo at least 3 passages to completely reprogram to a naïve-like state. At each passage, cells were dissociated using TrypLE (Thermo Fisher, 12604-021) and 10^6^ cells were seeded on a 10 cm culture dish with 5 mM ROCKi. ROCKi was withdrawn the day after thawing.

#### Naïve human PSC culture (PXGL)

H9 hESCs were cultured on gelatin-coated plates including a feeder layer of mitomycin treated mouse embryonic fibroblasts (MEFs) in PXGL medium. PXGL medium is prepared using N2B27 basal medium supplemented with PD0325901(1 µ M, MedChemExpress, HY-10254), XAV-939 (2 µM, MedChemExpress, HY-15147), Gö 6983 (2 µM, MedChemExpress, HY-13689) and human leukemia inhibitory factor (hLIF, 10 ng ml^−1^, in-house made). N2B27 basal medium contained DMEM/F12 (50%, GIBCO, 11320-074), neurobasal medium (50%, GIBCO, 21103-049), N-2 supplement (Thermo Fisher Science, 17502048), B-27 supplement (Thermo Fisher Science, 17504044), GultaMAX supplement (Thermo Fisher Science, 35050-038), non-essential amino acid, 2-mercaptoethanol (100 µM, Thermo Fisher Science, 31350010), and bovine serum albumin solution (0.45%, Sigma-Aldrich, A7979-50ML). Cells were routinely cultured in hypoxic chambers (5% CO_2_, 5% O_2_) and passaged as single cells every three to four days. All cell lines had routinely tested negative for mycoplasma.

#### Mouse ESC culture

Mouse ESCs were plated on 0.1% gelatin-coated dishes and grown in DMEM high glucose with Glutamax media (Thermo, 31966047) supplemented with 15% FBS (Thermo, 2206648RP), 1X NEAA (Thermo, 11140-035), 1X b-mercaptoethanol (Thermo, 21985023), 1X Penicillin/streptomycin (Life Technologies, 15140148) and 1000U/mL LIF and grown at 37°C in 20% O_2_ and 5% CO_2_ incubator. At each passage, cells were dissociated using TrypLE (Thermo Fisher, 12604-021) with media change every day.

### Developmental pausing setup

#### Human PSC pausing

For pausing of primed hiPSCs, cells were treated with the catalytic mTOR inhibitor INK128 (MedChemExpress/ Biozol, MCE-HY-13328) at 200 nM final concentration and 10 mM ROCKi in mTeSR media for 1 day. The following day, ROCKi was withdrawn and cells were cultured in media containing INK128 for six days with daily media change.

For pausing of hiPSCs in RSeT culture, 10^6^ cells were plated and grown in RSeT feeder-free medium. The media was changed every other day. On the fourth day of culture (approximate colony diameter of 100 mm), media containing 200 nM INK128 and 5 mM ROCKi was added for one day, then replaced with media containing only 200 nM of mTORi. Media was changed every day.

For pausing of hESCs in PXGL culture, 400,000 cells were counted and plated onto confluent MEFs in PXGL media with 10 mM ROCKi, Matrigel and 200 nM INK128 for one day. The next day the media was replenished with PXGL and 200 nM INK128. To validate INK128-mediated pausing, RapaLink-1 (Biozol, APE-A87764) was used at 200 nM and Torin1 (Abcam, ab218606) was used at 200-1000 nM.

#### Mouse ESC pausing

For pausing of mouse ESCs, cells were treated with mTOR inhibitor at 200nM final concentration and the cells were cultured for six days. Media was replenished as required.

#### Mouse blastocyst pausing

For embryo collection, F1 (C57Bl/6xCBA) females were superovulated to obtain fertilized oocytes. Superovulated female mice were set up for mating with eight-week-old or older (C57Bl/6xCBA) F1 male mice. E0.5 embryos were collected from swollen ampulas in FHM medium (Millipore #MR-024-D;27:50), treated with hyaluronidase (Sigma-Aldrich; H4272) to remove cumulus cells, and embryos were cultured in drops of pre-equilibrated KSOM medium overlaid with mineral oil (Origio; ART-4008-5P) at 37.5°C in 5% CO2. In vivo diapause was induced after natural mating of CD1 mice. Pregnant females were ovariectomized at E3.5 and afterwards injected every other day with 3 mg medroxyprogesterone 17-acetate (subcutaneously). Diapaused blastocysts were flushed from uteri in M2 media after 3 of diapause at EDG7.5.

### Immunofluorescence (IF)

#### PSCs

Cells were cultured on glass coverslips and were fixed in 4% PFA for 10 min at room temperature, washed once in PBS, then permeabilized with 0.2% Triton-X100 in PBS for 5 min on ice. After washing once in PBS-T (PBS with 0.2% Tween-20), cells were blocked with blocking buffer (PBS-T, 2% BSA and 5% goat serum (Jackson Immunoresearch/Dianova, 017-000-121) for 1 hour at room temperature. Cells were then stained with primary antibodies pS6 (CST Cat no: 4858) 1:200, pAKT (CST, 4060T) 1:200, KI67 (BD Pharmingen, 556003) 1:400, H3 phosphoS10 (Abcam, ab5176), 1:1000, OCT4 (Santa Cruz, sc5279) 1:50, NANOG (Abcam, ab109250) 1:200, gH2A.X (Biolegend, 613405) 1:400 overnight at 4°C. The cells were washed thrice with wash buffer (PBS-T, 2% BSA) for 10 min. Anti-rabbit (Thermo, A10042) or anti-mouse (Thermo, 21202) secondary antibody conjugated with Alexa Fluor was added to cells at a dilution of 1:700 in blocking buffer and incubated for 1 hour at room temperature, followed by 3 washes with wash buffer for 10 min. The coverslips were then mounted with Vectashield with DAPI (Vector labs, H-2000) and sealed with nail polish. Imaging was done using a Zeiss LSM880 Airy microscope using Airy scan mode and image processing was done using Zen black and Zen blue software (version 2.3). Image quantification was done using CellProfiler (version 4.2.1) where nuclei or cells which were denoted as primary objects were identified and the normalized intensities of the respective protein stained were measured against nuclear or cell area(https://cellprofiler.org/). Data were plotted using GraphPad Prism (version 9).

#### Embryos

Human embryos were fixed in 4% paraformaldehyde in PBS for 1h at 4°C, and mouse embryos 10 min at room temperature. Embryos were washed once in PBS, then permeabilized with 1× PBS with 0.5% Triton X-100 and then blocked in blocking solution (10% FBS in 1× PBS with 0.1% Triton X-100) for 1-2h at room temperature on a rotating shaker. Embryos were then incubated with primary antibodies diluted in blocking solution overnight at 4 °C on rotating shaker. The following day, embryos were washed once in 1× PBS with 0.1% Triton X-100 at room temperature on a rotating shaker, and then incubated with secondary antibodies diluted in blocking solution for 1 h at room temperature on a rotating shaker in the dark. Embryos were washed in 1× PBS with 0.1% Triton X-100 and counterstained with DAPI. The following antibodies and dilutions were used: anti-pS6 (Cell Signaling, 4858) 1:250, anti-OCT4 (Santa Cruz, 5279) 1:50, anti-Nanog (R&D AF1997) 1:200, anti-CDX2 (BioGenex, MU392-UC) 1:100, anti-pAKT (CST 4060T) 1:100, anti-4EBP1 (CST 2855) 1:100, anti-Lamin B1 (Abcam ab16048) 1:100, and anti-GATA3 (R&D AF2605) 1:200. All secondary antibodies were Alexa Fluor (Life Technologies), raised in donkey and used 1:200 to 1:1000. For imaging, embryos were placed on a µ-Slide 18 Well Flat dish (Ibidi, 81826) in PBS and imaged on a Leica Sp8 confocal with a Leica HCX PL APO 63x / 1.3 GLYC CORR CS objective. The z-stack step is 3.5-5 mm. Imaging of mouse embryos was done using a Zeiss LSM880 Airy microscope using Airy scan mode.

### Apoptosis assay

Cells adherent to the plate as well as floating cells were collected for the apoptosis assay. Cells were dissociated using TrypLE, washed in cold PBS, and resuspended in Annexin binding buffer (10 mM HEPES, 140 mM NaCl and 2.5 mM CaCl_2_, pH 7.4). Cell density was adjusted to 400,000 cells in 500 ml Annexin binding buffer. Staining for Annexin V was done according to manufacturer’s instructions (Thermo Fisher, R37174) along with dead cell stain using SYTOX AADvanced (Thermo, S10274) for 15 min at room temperature. A FACS AriaFusion flow cell cytometer was used to analyze cell staining. Data were analyzed using FlowJo (version 10) and plotted using GraphPad Prism (version 9).

### Cell cycle assay

To study cell cycle distribution, the Click-iT EdU Alexa Fluor 488 Flow Cytometry Assay kit (Thermo Fisher, C10425) was used. Normal and paused iPSCs were incubated at 37°C for 2 hours with 10 µM EdU in 5% O_2_, 5% CO_2_. Cells were then harvested, washed once with 3 ml of 1% BSA in PBS, centrifuged at 300 g for 5 min and the supernatant was removed. The pellet was dislodged and fixed in 100ml Click-iT fixative for 15 minutes at room temperature. After fixation, cells were washed with 3 ml of 1% BSA in PBS. The pellet was then resuspended in 100 ml 1x Click-iT saponin-based permeabilization and wash reagent and incubated for 15 minutes at room temperature. To this, 500 ml Click-iT reaction cocktail was added and incubated for 30 minutes in the dark at room temperature. After incubation, cells were washed with 3 ml of 1X Click-iT saponin based permeabilization and wash reagent, suspended in 200 ul of the same reagent. FxCycle violet (Life Technologies, F10347) was added to a final concentration of 1 mg/ml to measure DNA content and incubated for 1 hour at room temperature in the dark. FACS AriaFusion cell cytometer was used to acquire data using BD FACSDIVA Software v8.0.1. Data were analyzed using FlowJo (version 10) and plotted using GraphPad Prism (version 9).

### Global proteomics

#### Low-input proteomics

5000 cells per sample were lysed in a denaturing buffer, reduced and alkylated, and sequentially digested by Lys-C and trypsin. Peptides originating from about 1000 cells were loaded onto Evotips Pure (Evosep, Odense, Denmark) according to manufacturer protocol. Peptide separation was carried out by nanoflow reverse phase liquid chromatography (Evosep One, Evosep), using the Endurance column (15 cm x 150 µm ID, with Reprosil-Pur C18 1.9 µm beads #EV1106, Evosep) with the 30 samples a day method (30SPD). The LC system was online coupled to a timsTOF SCP mass spectrometer (Bruker Daltonics, Bremen, Germany) applying the data-independent acquisition (DIA) with parallel accumulation serial fragmentation (PASEF) method(*2*). MS data were processed with DiaNN (v1.8) and searched against in silico predicted mouse or human spectra (*3*).

#### Standard bulk proteomics

Proteomics sample preparation was done according to a published protocol with minor modifications(*4*). In brief, 5×10^6^ cells in biological duplicates were lysed under denaturing conditions in 500 µl of a buffer containing 3 M guanidinium chloride (Gdm-Cl), 10 mM tris(2-carboxyethyl)phosphine, 40 mM chloroacetamide, and 100 mM Tris-HCl pH 8.5. Lysates were denatured at 95°C for 10 min shaking at 1000 rpm in a thermal shaker and sonicated in a water bath for 10 min. 100 µl lysate was diluted with a dilution buffer containing 10% acetonitrile and 25 mM Tris-HCl, pH 8.0, to reach a 1 M GdmCl concentration. Then, proteins were digested with LysC (Roche, Basel, Switzerland; enzyme to protein ratio 1:50, MS-grade) shaking at 700 rpm at 37°C for 2 hours. The digestion mixture was diluted again with the same dilution buffer to reach 0.5 M GdmCl, followed by a tryptic digestion (Roche, enzyme to protein ratio 1:50, MS-grade) and incubation at 37°C overnight in a thermal shaker at 700 rpm. Peptide desalting was performed according to the manufacturer’s instructions (Pierce C18 Tips, Thermo Scientific, Waltham, MA). Desalted peptides were reconstituted in 0.1% formic acid in water and further separated into four fractions by strong cation exchange chromatography (SCX, 3M Purification, Meriden, CT). Eluates were first dried in a SpeedVac, then dissolved in 5% acetonitrile and 2% formic acid in water, briefly vortexed, and sonicated in a water bath for 30 seconds prior injection to nano-LC-MS/MS.

LC-MS/MS was carried out by nanoflow reverse phase liquid chromatography (Dionex Ultimate 3000, Thermo Scientific) coupled online to a Q-Exactive HF Orbitrap mass spectrometer (Thermo Scientific), as reported previously(*5*). Briefly, the LC separation was performed using a PicoFrit analytical column (75 µm ID × 50 cm long, 15 µm Tip ID; New Objectives, Woburn, MA) in-house packed with 3-µm C18 resin (Reprosil-AQ Pur, Dr. Maisch, Ammerbuch, Germany).

Raw MS data were processed with MaxQuant software (v1.6.10.43) and searched against the mouse proteome database UniProtKB with 55,153 entries, released in August 2019. The MaxQuant processed output files can be found in Table S1, showing peptide and protein identification, accession numbers, % sequence coverage of the protein, q-values, and label free quantification (LFQ) intensities.

#### Differential expression analysis

The Differential Enrichment analysis of Proteomics data (DEP) package v1.16.0 was used in R for proteomics data preparation and the statistical analysis(*6*). For human iPSCs, the label free quantification (LFQ) values were filtered. Only proteins quantified in both replicates of at least one condition were kept. Number of proteins kept after filtering: 4470 in PXGL, 4615 in RSeT, 5872 in primed conditions. For mouse ESCs, only proteins quantified in at least 2 out of 3 replicates of at least one condition were kept. A total of 4,783 proteins were kept after filtering. Both human and mouse proteomics data was background-corrected and normalized by variance stabilizing transformation. Missing values were imputed using random draws from a Gaussian distribution centered around a minimal value. Proteins with a p.adj < 0.05 and |log_2_FC| > 1 were considered differentially expressed. The differential protein expression analysis results of the human PSCs and mouse ESCs can be found in Table S2.

#### Scatter plots

The global change in proteome profile was displayed by the mean LFQ values in normal vs paused iPSCs and were generated using ggplot2.

#### GO term analysis

To identify enriched Biological Processes, Gene Ontology analysis in the clusterProfiler R package was applied on the differentially expressed proteins (DEPs) with a p.adj < 0.05 and |log2FC| of >1(*7*). The Benjamini-Hochberg correction was used to correct for multiple comparisons and a pvalueCutoff of 0.05 and qvalueCutoff of 0.1 were used. Enriched biological processes were displayed with a cnetplot and dotplot. Selected biological processes were displayed with ggplot2. A full overview of the enriched biological processes is provided in Table S3.

### Global proteomics: human-mouse comparison

#### Pairwise protein expression analysis

A total of 2861 proteins were expressed in both human (PXGL) and mouse data. The log_2_FC of paused vs proliferating of all overlapping proteins was plotted with ggscatter and the Spearman’s Rho correlation coefficient was calculated.

#### Pairwise pathway expression analysis

KEGG (*8*) pathways containing at least 10 genes symbols were included in the pairwise pathway expression analysis. A total of 146 pathways were shared between mouse and human data. The pathway expression value was defined as the mean log_2_FC of proteins between paused and proliferating mouse ESCs and human naïve iPSCs, or between different culture conditions for human PSCs. Pathway and gene log_2_FC for human and mouse are provided in Table S4. The mean log_2_FC for each pathway for human and mouse data was plotted with ggscatter and the Spearman’s Rho correlation coefficient was calculated.

##### Pseudotime analysis

Data was filtered with only those proteins remaining that were expressed in both samples of at least one condition. Data was normalised and missing values were imputed using random draws from a Gaussian distribution centred around a minimal value with the DEP package. Proteins expressed in both the human and mouse ES cells were used to compute a diffusion map pseudotime with the destiny package (v3.10) in R (version 4.2.1). The species-specific data was used to generate a pseudotime based on the ranked diffusion component, using destiny package.

##### Human blastoid formation

Naive hPSCs were cultured under humidified conditions at 37°C in an incubator with 5% O_2_ and 5% CO_2_. Naive H9 hPSCs were cultured on mitotically inactivated mouse embryonic fibroblasts (MEFs) in PXGL medium. The medium was changed every day. The cells were passaged 4 d before blastoid formation. Blastoids were formed as previously published with minor modification (*9, 10*). After 4 days, blastoids were treated with 100, 200 and 300 nM RapaLink-1 (*13*), 200 nM INK128, 100 and 1000 ng/ml cycloheximide. The phase-contrast images were acquired using Thermo Fisher scientific EVOS cell imaging system. The number of blastoids in the microwells were counted manually for each well every day. For the survival curves, only visible blastoids in the microwells were counted.

### Blastoid reactivation after pausing

Paused blastoids were reactivated by culturing them in an extended culture condition previously published for human blastoids (*10*). Blastoids were selected using a mouth pipette, washed with CMRL1066 medium and transferred into wells of a 96-well plate coated with Matrigel containing pre-equilibrated media. For the first day, the culture medium was CMRL1066 (*14*) supplemented with 10% (v/v) FBS, 1 mM L-glutamine (Gibco), 1× N2 supplement, 1× B27 supplement, 1 mM sodium pyruvate (Sigma) and 10 µM Y27632. After 24 h, half of the medium was replaced with a new medium including 5% Matrigel. After 48h, 50% of medium was replaced with a new medium supplemented with 20% (v/v) FBS and 5% Matrigel. After 72h, half of the medium was replaced with a new medium supplemented with 30% (v/v) KSR and 5% Matrigel. Cultures were fixed for staining after 2 or 4 days of culture with 4% PFA.

### Immunofluorescence microscopy

Blastoids were collected using mouth pipette and transferred to U-bottomed 96-well plates (Merck, BR701330). Once the structures had settled, the medium was washed twice with PBS and fixed with 4% PFA for 30 min at room temperature, followed by three 10 min washes with PBS. PBS containing 10% normal donkey serum and 0.3% Triton X-100 was used for blocking and permeabilization for 3h at room temperature. The primary antibody was incubated at 4 °C in blocking/permeabilization solution with gentle shaking and washed at least three times with PBS containing 0.1% Triton X-100 for 10 min. The secondary antibody was diluted in PBS containing 0.1% Triton X-100 and incubated at room temperature in the dark for 1 h. Then, the blastoids were washed three times with PBS containing 0.1% Triton X-100 for 10 min and prepared for imaging. For imaging, the blastoids were placed in a glass-bottomed plates. Confocal IF images of blastoids were acquired with an Olympus IX83 microscope with Yokogawa W1 spinning disk (Software: CellSense 2.3; camera: Hamamatsu Orca Flash 4.0). The confocal images were analyzed, and display images were exported using FIJI 1.53k or Bitplane IMARIS 9.7.0 software. For cell counting, Bitplane IMARIS software was used. Cell count parameters were set for size and fluorescence strength of voxels and then overall cell count data was obtained for each image using the IMARIS spot function.

## REFERENCES

1. A. Bulut-Karslioglu, S. Biechele, H. Jin, T. A. Macrae, M. Hejna, M. Gertsenstein, J. S. Song, M. Ramalho-Santos, Inhibition of mTOR induces a paused pluripotent state. Nature. 540, 119–123 (2016).

2. M. B. Renfree, J. C. Fenelon, The enigma of embryonic diapause. Development. 144, 3199–3210 (2017).

3. V. A. van der Weijden, A. Bulut-Karslioglu, Molecular Regulation of Paused Pluripotency in Early Mammalian Embryos and Stem Cells. Frontiers Cell Dev Biology. 9, 708318 (2021).

4. V. A. van der Weijden, S. E. Ulbrich, Embryonic diapause in roe deer: A model to unravel embryomaternal communication during pre-implantation development in wildlife and livestock species. Theriogenology. 158, 105–111 (2020).

5. T. Boroviak, R. Loos, P. Lombard, J. Okahara, R. Behr, E. Sasaki, J. Nichols, A. Smith, P. Bertone, Lineage-Specific Profiling Delineates the Emergence and Progression of Naive Pluripotency in Mammalian Embryogenesis. Developmental Cell. 35, 366–382 (2015).

6. G. E. Ptak, E. Tacconi, M. Czernik, P. Toschi, J. A. Modlinski, P. Loi, Embryonic diapause is conserved across mammals. PloS one. 7, e33027 (2012).

7. M. C. Chang, Reciprocal insemination and egg transfer between ferrets and mink. J Exp Zool. 168, 49–59 (1968).

8. R. Alberio, T. Kobayashi, M. A. Surani, Conserved features of non-primate bilaminar disc embryos and the germline. Stem Cell Rep. 16, 1078–1092 (2021).

9. T. Rayon, D. Stamataki, R. Perez-Carrasco, L. Garcia-Perez, C. Barrington, M. Melchionda, K. Exelby, J. Lazaro, V. L. J. Tybulewicz, E. M. C. Fisher, J. Briscoe, Species-specific pace of development is associated with differences in protein stability. Science. 369, eaba7667 (2020).

10. M. Roode, K. Blair, P. Snell, K. Elder, S. Marchant, A. Smith, J. Nichols, Human hypoblast formation is not dependent on FGF signalling. Dev Biol. 361, 358–363 (2012).

11. M. Zhu, M. Shahbazi, A. Martin, C. Zhang, B. Sozen, M. Borsos, R. S. Mandelbaum, R. J. Paulson, M. A. Mole, M. Esbert, S. Titus, R. T. Scott, A. Campbell, S. Fishel, V. Gradinaru, H. Zhao, K. Wu, Z.-J. Chen, E. Seli, M. J. de los Santos, M. Z. Goetz, Human embryo polarization requires PLC signaling to mediate trophectoderm specification. Elife. 10, e65068 (2021).

12. E. W. Kuijk, L. T. A. van Tol, H. V. de Velde, R. Wubbolts, M. Welling, N. Geijsen, B. A. J. Roelen, The roles of FGF and MAP kinase signaling in the segregation of the epiblast and hypoblast cell lineages in bovine and human embryos. Development. 139, 871–882 (2012).

13. C. Gerri, A. McCarthy, G. Alanis-Lobato, A. Demtschenko, A. Bruneau, S. Loubersac, N. M. E. Fogarty, D. Hampshire, K. Elder, P. Snell, L. Christie, L. David, H. V. de Velde, A. A. Fouladi-Nashta, K. K. Niakan, Initiation of a conserved trophectoderm program in human, cow and mouse embryos. Nature. 587, 443–447 (2020).

14. L. Yan, M. Yang, H. Guo, L. Yang, J. Wu, R. Li, P. Liu, Y. Lian, X. Zheng, J. Yan, J. Huang, M. Li, X. Wu, L. Wen, K. Lao, R. Li, J. Qiao, F. Tang, Single-cell RNA-Seq profiling of human preimplantation embryos and embryonic stem cells. Nature Structural & Molecular Biology. 20, 1131–1139 (2013).

15. S. Petropoulos, D. Edsgärd, B. Reinius, Q. Deng, S. P. Panula, S. Codeluppi, A. Plaza Reyes, S. Linnarsson, R. Sandberg, F. Lanner, Single-Cell RNA-Seq Reveals Lineage and X Chromosome Dynamics in Human Preimplantation Embryos. Cell. 165, 1012–1026 (2016).

16. G. E. Ptak, J. A. Modlinski, P. Loi, Embryonic diapause in humans: time to consider? Reproductive biology and endocrinology: RB&E. 11, 92 (2013).

17. J. Grinsted, B. Avery, A sporadic case of delayed implantation after in-vitro fertilization in the human? Hum Reprod. 11, 651–654 (1996).

18. W. M. Liu, R. R. Cheng, Z. R. Niu, A. C. Chen, M. Y. Ma, T. Li, P. C. Chiu, R. T. Pang, Y. L. Lee, J. P. Ou, Y. Q. Yao, W. S. B. Yeung, Let-7 derived from endometrial extracellular vesicles is an important inducer of embryonic diapause in mice. Sci Adv. 6, eaaz7070 (2020).

19. I. Canton, N. J. Warren, A. Chahal, K. Amps, A. Wood, R. Weightman, E. Wang, H. Moore, S. P. Armes, Mucin-Inspired Thermoresponsive Synthetic Hydrogels Induce Stasis in Human Pluripotent Stem Cells and Human Embryos. Acs Central Sci. 2, 65– 74 (2016).

20. V. A. van der Weijden, M. Stoetzel, B. Fauler, D. P. Iyer, M. Shahraz, D. Meierhofer, S. Rulands, T. Alexandrov, T. Mielke, A. Bulut-Karslioglu, Metabolic enhancement of mammalian developmental pausing (2022), doi:10.1101/2022.08.22.504730.

21. C. Kamemizu, T. Fujimori, Distinct dormancy progression depending on embryonic regions during mouse embryonic diapause†. Biol Reprod. 100, 1204–1214 (2018).

22. G. Y. Liu, D. M. Sabatini, mTOR at the nexus of nutrition, growth, ageing and disease. Nat Rev Mol Cell Bio. 21, 183–203 (2020).

23. S. C. Woll, J. E. Podrabsky, Insulin-like growth factor signaling regulates developmental trajectory associated with diapause in embryos of the annual killifish Austrofundulus limnaeus. J Exp Biol. 220, 2777–2786 (2017).

24. K. D. Kimura, H. A. Tissenbaum, Y. Liu, G. Ruvkun, daf-2, an Insulin Receptor-Like Gene That Regulates Longevity and Diapause in Caenorhabditis elegans. Science. 277, 942–946 (1997).

25. P. Ciarmela, Md. S. Islam, F. M. Reis, P. C. Gray, E. Bloise, F. Petraglia, W. Vale, M. Castellucci, Growth factors and myometrium: biological effects in uterine fibroid and possible clinical implications. Hum Reprod Update. 17, 772–790 (2011).

26. L. Peng, Y. Wen, Y. Han, A. Wei, G. Shi, M. Mizuguchi, P. Lee, E. Hernando, K. Mittal, J.-J. Wei, Expression of insulin-like growth factors (IGFs) and IGF signaling: molecular complexity in uterine leiomyomas. Fertil Steril. 91, 2664–2675 (2009).

27. A. D. Lighten, G. E. Moore, R. M. Winston, K. Hardy, Routine addition of human insulin-like growth factor-I ligand could benefit clinical in-vitro fertilization culture. Hum Reprod. 13, 3144–3150 (1998).

28. H. Kagawa, A. Javali, H. H. Khoei, T. M. Sommer, G. Sestini, M. Novatchkova, Y. S. op Reimer, G. Castel, A. Bruneau, N. Maenhoudt, J. Lammers, S. Loubersac, T. Freour, H. Vankelecom, L. David, N. Rivron, Human blastoids model blastocyst development and implantation. Nature. 601, 600–605 (2022).

29. C. Zhao, A. P. Reyes, J. P. Schell, J. Weltner, N. M. Ortega, Y. Zheng, Å. K. Björklund, J. Rossant, J. Fu, S. Petropoulos, F. Lanner, Biorxiv, in press, doi:10.1101/2021.05.07.442980.

30. J. Nichols, I. Chambers, T. Taga, A. Smith, Physiological rationale for responsiveness of mouse embryonic stem cells to gp130 cytokines. Development. 128, 2333–2339 (2001).

31. T. W. Theunissen, B. E. Powell, H. Wang, M. Mitalipova, D. A. Faddah, J. Reddy, Z. P. Fan, D. Maetzel, K. Ganz, L. Shi, T. Lungjangwa, S. Imsoonthornruksa, Y. Stelzer, S. Rangarajan, A. D’Alessio, J. Zhang, Q. Gao, M. M. Dawlaty, R. A. Young, N. S. Gray, R. Jaenisch, Systematic Identification of Culture Conditions for Induction and Maintenance of Naive Human Pluripotency. Cell Stem Cell. 15, 471–487 (2014).

32. Y. Takashima, G. Guo, R. Loos, J. Nichols, G. Ficz, F. Krueger, D. Oxley, F. Santos, J. Clarke, W. Mansfield, W. Reik, P. Bertone, A. Smith, Resetting Transcription Factor Control Circuitry toward Ground-State Pluripotency in Human. Cell. 158, 1254–1269 (2014).

33. Y.-S. Chan, J. Göke, J.-H. Ng, X. Lu, K. A. U. Gonzales, C.-P. Tan, W.-Q. Tng, Z.-Z. Hong, Y.-S. Lim, H.-H. Ng, Induction of a Human Pluripotent State with Distinct Regulatory Circuitry that Resembles Preimplantation Epiblast. Cell Stem Cell. 13, 663–675 (2013).

34. G. Guo, F. von Meyenn, F. Santos, Y. Chen, W. Reik, P. Bertone, A. Smith, J. Nichols, Naive Pluripotent Stem Cells Derived Directly from Isolated Cells of the Human Inner Cell Mass. Stem Cell Reports. 6, 437–446 (2016).

35. O. Gafni, L. Weinberger, A. A. Mansour, Y. S. Manor, E. Chomsky, D. Ben-Yosef, Y. Kalma, S. Viukov, I. Maza, A. Zviran, Y. Rais, Z. Shipony, Z. Mukamel, V. Krupalnik, M. Zerbib, S. Geula, I. Caspi, D. Schneir, T. Shwartz, S. Gilad, D. Amann-Zalcenstein, S. Benjamin, I. Amit, A. Tanay, R. Massarwa, N. Novershtern, J. H. Hanna, Derivation of novel human ground state naive pluripotent stem cells. Nature. 504, 282–286 (2013).

36. C. B. Ware, A. M. Nelson, B. Mecham, J. Hesson, W. Zhou, E. C. Jonlin, A. J. Jimenez-Caliani, X. Deng, C. Cavanaugh, S. Cook, P. J. Tesar, J. Okada, L. Margaretha, H. Sperber, M. Choi, C. A. Blau, P. M. Treuting, R. D. Hawkins, V. Cirulli, H. Ruohola-Baker, Derivation of naïve human embryonic stem cells. Proc National Acad Sci. 111, 4484–4489 (2014).

37. S. Giulitti, M. Pellegrini, I. Zorzan, P. Martini, O. Gagliano, M. Mutarelli, M. J. Ziller, D. Cacchiarelli, C. Romualdi, N. Elvassore, G. Martello, Direct generation of human naive induced pluripotent stem cells from somatic cells in microfluidics. Nat Cell Biol. 21, 275–286 (2019).

38. J. Bayerl, M. Ayyash, T. Shani, Y. S. Manor, O. Gafni, R. Massarwa, Y. Kalma, A. Aguilera-Castrejon, M. Zerbib, H. Amir, D. Sheban, S. Geula, N. Mor, L. Weinberger, S. N. Tassa, V. Krupalnik, B. Oldak, N. Livnat, S. Tarazi, S. Tawil, E. Wildschutz, S. Ashouokhi, L. Lasman, V. Rotter, S. Hanna, D. Ben-Yosef, N. Novershtern, S. Viukov, J. H. Hanna, Principles of signaling pathway modulation for enhancing human naive pluripotency induction. Cell Stem Cell. 28, 1549–1565.e12 (2021).

39. S. Kilens, D. Meistermann, D. Moreno, C. Chariau, A. Gaignerie, A. Reignier, Y. Lelièvre, M. Casanova, C. Vallot, S. Nedellec, L. Flippe, J. Firmin, J. Song, E. Charpentier, J. Lammers, A. Donnart, N. Marec, W. Deb, A. Bihouée, C. L. Caignec, C. Pecqueur, R. Redon, P. Barrière, J. Bourdon, V. Pasque, M. Soumillon, T. S. Mikkelsen, C. Rougeulle, T. Fréour, L. David, L. Abel, A. Alcover, K. Astrom, P. Bousso, P. Bruhns, A. Cumano, D. Duffy, C. Demangel, L. Deriano, J. D. Santo, F. Dromer, G. Eberl, J. Enninga, J. Fellay, A. Freitas, O. Gelpi, I. Gomperts-Boneca, S. Hercberg, O. Lantz, C. Leclerc, H. Mouquet, E. Patin, S. Pellegrini, S. Pol, L. Rogge, A. Sakuntabhai, O. Schwartz, B. Schwikowski, S. Shorte, V. Soumelis, F. Tangy, E. Tartour, A. Toubert, M.-N. Ungeheuer, L. Quintana-Murci, M. L. Albert, Parallel derivation of isogenic human primed and naive induced pluripotent stem cells. Nat Commun. 9, 360 (2018).

40. Z. Hu, H. Li, H. Jiang, Y. Ren, X. Yu, J. Qiu, A. B. Stablewski, B. Zhang, M. J. Buck, J. Feng, Transient inhibition of mTOR in human pluripotent stem cells enables robust formation of mouse-human chimeric embryos. Sci Adv. 6, eaaz0298 (2020).

41. H. Qin, M. Hejna, Y. Liu, M. Percharde, M. Wossidlo, L. Blouin, J. Durruthy-Durruthy, P. Wong, Z. Qi, J. Yu, L. S. Qi, V. Sebastiano, J. S. Song, M. Ramalho-Santos, YAP Induces Human Naive Pluripotency. Cell reports. 14, 2301–2312 (2016).

42. L. Weinberger, M. Ayyash, N. Novershtern, J. H. Hanna, Dynamic stem cell states: naive to primed pluripotency in rodents and humans. Nat Rev Mol Cell Bio. 17, 155–169 (2016).

43. R. Arena, S. Bisogno, Ł. Gąsior, J. Rudnicka, L. Bernhardt, T. Haaf, F. Zacchini, M. Bochenek, K. Fic, E. Bik, M. Barańska, A. Bodzoń-Kułakowska, P. Suder, J. Depciuch, A. Gurgul, Z. Polański, G. E. Ptak, Lipid droplets in mammalian eggs are utilized during embryonic diapause. Proc National Acad Sci. 118, e2018362118 (2021).

44. E. S. Sills, X. Li, J. L. Frederick, C. D. Khoury, D. A. Potter, Determining parental origin of embryo aneuploidy: analysis of genetic error observed in 305 embryos derived from anonymous donor oocyte IVF cycles. Mol Cytogenet. 7, 68 (2014).

45. M. J. Zinaman, E. D. Clegg, C. C. Brown, J. O’-Connor, S. G. Selevan, Estimates of human fertility and pregnancy loss. Fertil Steril. 65, 503–509 (1996).

46. T.-C. Lin, J.-M. Yen, K.-B. Gong, T.-T. Hsu, L.-R. Chen, IGF-1/IGFBP-1 increases blastocyst formation and total blastocyst cell number in mouse embryo culture and facilitates the establishment of a stem-cell line. Bmc Cell Biol. 4, 14 (2003).

47. S. E. Wamaitha, K. J. Grybel, G. Alanis-Lobato, C. Gerri, S. Ogushi, A. McCarthy, S. K. Mahadevaiah, L. Healy, R. A. Lea, M. Molina-Arcas, L. G. Devito, K. Elder, P. Snell, L. Christie, J. Downward, J. M. A. Turner, K. K. Niakan, IGF1-mediated human embryonic stem cell self-renewal recapitulates the embryonic niche. Nat Commun. 11, 764 (2020).

48. J. K. Riley, K. H. Moley, Glucose utilization and the PI3-K pathway: mechanisms for cell survival in preimplantation embryos. Reproduction. 131, 823–835 (2006).

49. S. Pauklin, L. Vallier, The Cell-Cycle State of Stem Cells Determines Cell Fate Propensity. Cell. 155, 135–147 (2013).

50. J. A. Saba, K. Liakath-Ali, R. Green, F. M. Watt, Translational control of stem cell function. Nat Rev Mol Cell Bio. 22, 671–690 (2021).

51. M. Buszczak, R. A. J. Signer, S. J. Morrison, Cellular differences in protein synthesis regulate tissue homeostasis. Cell. 159, 242–251 (2014).

52. R. A. J. Signer, J. A. Magee, A. Salic, S. J. Morrison, Haematopoietic stem cells require a highly regulated protein synthesis rate. Nature. 509, 49–54 (2014).

53. A. Bulut-Karslioglu, T. A. Macrae, J. A. Oses-Prieto, S. Covarrubias, M. Percharde, G. Ku, A. Diaz, M. T. McManus, A. L. Burlingame, M. Ramalho-Santos, The Transcriptionally Permissive Chromatin State of Embryonic Stem Cells Is Acutely Tuned to Translational Output. Cell Stem Cell. 22, 369–383.e8 (2018).

54. L. Martens, H. Hermjakob, P. Jones, M. Adamski, C. Taylor, D. States, K. Gevaert, J. Vandekerckhove, R. Apweiler, PRIDE: The proteomics identifications database. Proteomics. 5, 3537–3545 (2005).

## References

1. A. T. Clark, A. Brivanlou, J. Fu, K. Kato, D. Mathews, K. K. Niakan, N. Rivron, M. Saitou, A. Surani, F. Tang, J. Rossant, Human embryo research, stem cell-derived embryo models and in vitro gametogenesis: Considerations leading to the revised ISSCR guidelines. Stem Cell Rep. 16, 1416–1424 (2021).

2. F. Meier, A.-D. Brunner, S. Koch, H. Koch, M. Lubeck, M. Krause, N. Goedecke, J. Decker, T. Kosinski, M. A. Park, N. Bache, O. Hoerning, J. Cox, O. Räther, M. Mann, Online Parallel Accumulation–Serial Fragmentation (PASEF) with a Novel Trapped Ion Mobility Mass Spectrometer*. Mol Cell Proteomics. 17, i–2545 (2018).

3. V. Demichev, L. Szyrwiel, F. Yu, G. C. Teo, G. Rosenberger, A. Niewienda, D. Ludwig, J. Decker, S. Kaspar-Schoenefeld, K. S. Lilley, M. Mülleder, A. I. Nesvizhskii, M. Ralser, dia-PASEF data analysis using FragPipe and DIA-NN for deep proteomics of low sample amounts. Nat Commun. 13, 3944 (2022).

4. N. A. Kulak, G. Pichler, I. Paron, N. Nagaraj, M. Mann, Minimal, encapsulated proteomic-sample processing applied to copy-number estimation in eukaryotic cells. Nat Methods. 11, 319–324 (2014).

5. Y. Ni, M. A. Hagras, V. Konstantopoulou, J. A. Mayr, A. A. Stuchebrukhov, D. Meierhofer, Mutations in NDUFS1 Cause Metabolic Reprogramming and Disruption of the Electron Transfer. Cells. 8, 1149 (2019).

6. X. Zhang, A. H. Smits, G. B. van Tilburg, H. Ovaa, W. Huber, M. Vermeulen, Proteome-wide identification of ubiquitin interactions using UbIAMS. Nat Protoc. 13, 530–550 (2018).

7. T. Wu, E. Hu, S. Xu, M. Chen, P. Guo, Z. Dai, T. Feng, L. Zhou, W. Tang, L. Zhan, X. Fu, S. Liu, X. Bo, G. Yu, clusterProfiler 4.0: A universal enrichment tool for interpreting omics data. Innovation. 2, 100141 (2021).

8. M. Kanehisa, S. Goto, KEGG: Kyoto Encyclopedia of Genes and Genomes. Nucleic Acids Res. 28, 27–30 (2000).

9. H. Kagawa, A. Javali, H. H. Khoei, T. M. Sommer, G. Sestini, M. Novatchkova, Y. S. op Reimer, N. Rivron, Protocol for Human Blastoids Modeling Blastocyst Development and Implantation. J Vis Exp (2022), doi:10.3791/63388.

10. H. Kagawa, A. Javali, H. H. Khoei, T. M. Sommer, G. Sestini, M. Novatchkova, Y. S. op Reimer, G. Castel, A. Bruneau, N. Maenhoudt, J. Lammers, S. Loubersac, T. Freour, H. Vankelecom, L. David, N. Rivron, Human blastoids model blastocyst development and implantation. Nature. 601, 600–605 (2022).

11. E. J. Vrij, S. Espinoza, M. Heilig, A. Kolew, M. Schneider, C. A. van Blitterswijk, R. K. Truckenmüller, N. C. Rivron, 3D high throughput screening and profiling of embryoid bodies in thermoformed microwell plates. Lab Chip. 16, 734–742 (2016).

12. N. C. Rivron, E. J. Vrij, J. Rouwkema, S. L. Gac, A. van den Berg, R. K. Truckenmüller, C. A. van Blitterswijk, Tissue deformation spatially modulates VEGF signaling and angiogenesis. Proc National Acad Sci. 109, 6886–6891 (2012).

13. V. S. Rodrik-Outmezguine, M. Okaniwa, Z. Yao, C. J. Novotny, C. McWhirter, A. Banaji, H. Won, W. Wong, M. Berger, E. de Stanchina, D. G. Barratt, S. Cosulich, T. Klinowska, N. Rosen, K. M. Shokat, Overcoming mTOR resistance mutations with a new-generation mTOR inhibitor. Nature. 534, 272–276 (2016).

14. H. Ma, J. Zhai, H. Wan, X. Jiang, X. Wang, L. Wang, Y. Xiang, X. He, Z.-A. Zhao, B. Zhao, P. Zheng, L. Li, H. Wang, In vitro culture of cynomolgus monkey embryos beyond early gastrulation. Science. 366 (2019), doi:10.1126/science.aax7890.

